# O-GlcNAcylation of the intellectual disability protein DDX3X exerts proteostatic cell cycle control

**DOI:** 10.1101/2024.02.28.582457

**Authors:** Conor W. Mitchell, Huijie Yuan, Marie Sønderstrup-Jensen, Florence Authier, Alfonso Manuel D’Alessio, Andrew T. Ferenbach, Daan M.F. van Aalten

## Abstract

O-GlcNAcylation is an evolutionary conserved post-translational modification implicated in neurodevelopment. Missense variants of O-GlcNAc transferase (OGT) are causal for the intellectual disability syndrome OGT Congenital Disorder of Glycosylation (OGT-CDG). The observation of microcephaly in OGT-CDG patients suggests that dysregulation of the cell cycle and aberrant neurogenesis may contribute to disease aetiology. Here, we identify Ser584 O-GlcNAcylation of DDX3X, a known intellectual disability and microcephaly associated protein, as a key regulator of G1/S-phase transition, inhibiting proteasome-dependent degradation of DDX3X. DDX3X levels are reduced in a mouse model of OGT-CDG, alongside the DDX3X-target gene and synaptogenic regulator cyclin E1. These data reveal how a single DDX3X O-GlcNAc site exerts control of the cell cycle and highlights dysregulation of DDX3X-dependent translation, and concomitant impairments in cortical neurogenesis, as a possible pathway disrupted in OGT-CDG.

## Introduction

Modification of Ser/Thr hydroxyls with the monosaccharide O-linked-β-N-acetyl-glucosamine (O-GlcNAc) is a highly conserved and widespread post-translational modification (PTM), occurring on > 5000 nuclear, cytoplasmic, and mitochondrial proteins [1, 2]. O-GlcNAc is a functionally versatile PTM in cellular physiology, playing key roles in regulating gene expression programmes [3], proteostasis [4], transcriptional activation [4], and the stress response [5]. In contrast to other PTMs such as phosphorylation and ubiquitylation, where thousands of writers, readers, and erasers collectively generate and interpret the phospho- and ubiquitin proteomes [6, 7], O-GlcNAc homeostasis is regulated by the activity of just two, highly evolutionarily conserved, enzymes, the O-GlcNAc transferase (OGT) and O-GlcNAc hydrolase (OGA) [8, 9]. OGT is a member of the GT41 family of glycosyltransferases, with an N-terminal tetratricopeptide repeat (TPR) domain that is thought to be responsible for substrate binding and the binding of OGT interactors [10, 11]. Aside from catalysing O-GlcNAcylation, the C-terminal glycosyltransferase domain also catalyses proteolytic maturation of the transcription factor Host Cell Factor 1 (HCF1) in the same active site [12, 13].

*OGT* resides on the X-chromosome and is essential for development, as germline knockout of *Ogt* in mice is embryonic lethal [14]. This extends to the developing central nervous system (CNS), with conditional deletion of *Ogt* in neurons resulting in perinatal lethality accompanied with hallmarks of neurodegeneration [15]. Despite this barrier to dissecting the function of O-GlcNAc in neurodevelopment, several lines of evidence point to O-GlcNAc as a crucial signalling node for the development and functional maintenance of the CNS. Cell population specific knock-out of *Ogt* has previously demonstrated the importance of O-GlcNAcylation in dopaminergic neuronal survival, cortical synapse maturation, and regulation of satiety in the paraventricular nucleus of the hypothalamus [16–18]. Additionally, OGT and O-GlcNAc are enriched in the brain [3], particularly the hippocampus [4], with O-GlcNAcylation being abundant at neuronal synapses [4], and elevation of global O-GlcNAc levels through chemical and genetic methods inhibits long-term potentiation of hippocampal neurons, a key correlate of learning and memory [5]. Although the identity of the O-GlcNAc signalling nodes responsible for neuronal survival and synaptic transmission remain unknown, O-GlcNAc proteomic studies have estimated that approximately 20% of all synaptosomal proteins are O-GlcNAc modified, including proteins involved in signal transduction, neurite structure, and ion channels, providing insights into how O-GlcNAc signalling may modulate neurobiological outcomes [19]. To summarise, O-GlcNAcylation is a functionally diverse PTM implicated in the regulation of a plethora of cellular and developmental processes, particularly in the developing and post-natal nervous system.

Recently, missense variants in both the N-terminal TPR and C-terminal glycosyltransferase domains of OGT have been reported to segregate with a novel form of X-linked intellectual disability – OGT Congenital Disorder of Glycosylation (OGT-CDG) [20–25]. Intellectual disabilities are a clinically and genetically heterogeneous collection of developmental disorders characterised by reduced intellectual functioning, accompanied with impaired social or adaptive behaviours, with onset prior to 18 years of age [26]. OGT-CDG patients, aside from intellectual, social, and behavioural impairments, present with a wide range of facial, muscular, and neurological abnormalities, with hypotonia, 5^th^-finger clinodactyly, as well as microcephaly and hypoplasia of the corpus callosum frequently reported [20–22]. However, additional lower frequency symptoms such as epilepsy and autistic features have been reported in OGT-CDG patients [24, 25], suggesting a degree of clinical heterogeneity.

Since the discovery of OGT-CDG variants, divergent hypotheses have emerged regarding the aetiology of this novel ID syndrome [27]. The TPR domain variants that have been characterised to date display variable effects on protein folding and catalytic activity, suggesting a mixture of dosage and catalytic effects on OGT activity [23–25]. Conversely, catalytic domain variants reported to date exclusively affect catalytic activity *in vitro* and in both *Drosophila melanogaster* and embryonic stem cell models, without impaired protein folding or structural alterations *in crystallo* [23–25]. The latter finding suggests that OGT-CDG may result from hypo-O-GlcNAcylation of key O-GlcNAc signalling nodes, either during the early stages of neurodevelopment *in utero*, or postnatally during neurogenesis and in post-mitotic neurons. However, the role of O-GlcNAc on the majority of OGT substrates is presently unknown, hampering identification and dissection of O-GlcNAc-dependent signalling pathways, and establishing how disruption of these pathways contributes OGT-CDG.

Microcephaly, a neurodevelopmental condition resulting in reduced head circumference and thinning of the cerebral cortex [28], has been reported for three different OGT-CDG variants to date [20–22]. Generally, microcephaly originates from the early stages of corticogenesis when the radial glial cells (RGCs) undergo both symmetric and asymmetric rounds of cell division to either maintain the pool of neural progenitors or generate intermediate progenitors and post-mitotic neurons [29]. The latter subsequently migrate away from the ventricular surface of the developing cortex to form the six different layers of the cerebral cortex [29]. Microcephaly is closely associated with impaired cell cycle progression and cell division in neural progenitors, with mutations in centrosomal proteins [30, 31], cell cycle regulators [32–34], and components of the DNA damage response pathway resulting in delayed cell cycle and mitotic progression [35, 36], neural progenitor apoptosis [37], and genomic instability of RGC daughter cells [30, 35]. These defects collectively result in depletion of the RGC pool and a decrease in the number of cortical neurons [31], driving subsequent microcephaly in patients. It is therefore possible that either impaired mitotic or cell cycle progression of neural progenitors could underpin the microcephaly observed in OGT-CDG patients.

The requirement for O-GlcNAcylation in both cell cycle progression and the faithful transmission of genetic material to daughter cells has been previously established through transfection experiments *in cellulo*. Over-expression of OGT results in the formation of multi-polar spindles and aneuploidy, potentially due to increased inhibitory phosphorylation of the M phase regulator CDK1 [38, 39], whereas chemical inhibition of OGT results in stalled S-phase entry through as yet unidentified mechanisms [38, 40]. Although the broad cytological effects of O-GlcNAc on cell cycle progression have been reported, targeted dissection of O-GlcNAc signalling nodes and their contribution to cell cycle regulation is lacking. Previous studies have established the stabilising effect of O-GlcNAc on β-catenin, a transcription factor required for *cyclin D* transcription and cell cycle re-entry of quiescent cells [41]. Moreover, cleavage of HCF1 by OGT produces two functionally distinct HCF1 fragments [12, 13], with the N- and C-terminal fragments required for G1 progression and mitotic exit respectively. Aside from these studies, however, limited mechanistic detail is available. Given the potential link between the cell cycle, O-GlcNAcylation, and microcephaly, further elucidation of the regulatory pathways under the control of O-GlcNAcylation and dissection of how disruption of these processes might lead to microcephaly could lead to an increase in understanding of the processes underpinning OGT-CDG.

Here, we describe how O-GlcNAcylation of the intellectual disability protein DDX3X exerts proteostatic cell cycle control. DDX3X is an RNA helicase of the DEAD box family, and promotes cell cycle progression through translational control of genes including *cyclin E1* [42, 43]. Missense and nonsense variants in *DDX3X* cause an ID syndrome that presents with microcephaly due to impaired neural progenitor proliferation during corticogenesis [32, 44, 45]. Analysis of available expression data revealed a strong correlation between *DDX3X*, *OGT* and *OGA* expression in the frontal lobe of the cerebral cortex, suggesting mechanistic interplay between protein O-GlcNAcylation and DDX3X. Further investigation *in cellulo* revealed that Ser584 O-GlcNAcylation prevents DDX3X degradation by the ubiquitin-proteasome system. Loss of Ser584 O-GlcNAcylation increased DDX3X turnover sufficiently to impair the G1/S transition of the cell cycle. In line with this observation, we report reduced levels of DDX3X and cyclin E1 in the brains of mice carrying an OGT-CDG mutation. These findings describe a novel mechanism by which a single O-GlcNAc site regulates S-phase entry through fine-tuning of an essential cell cycle protein, highlight the possible convergence of cell cycle dysregulation and OGT-CDG aetiology, and suggest DDX3X as a candidate conveyor of the microcephaly observed in OGT-CDG patients.

## Results

### Expression of cell cycle regulators and intellectual disability associated genes, including DDX3X, are correlated with OGT and OGA in the human prefrontal cortex

As a strategy towards identifying signaling nodes and pathways regulated by OGT/OGA as candidate conveyors of OGT-CDG, we analysed available temporal transcriptomics of the human prefrontal cortex, a critical brain structure for executive decision making and working memory. We focused on genes correlated with *OGT* and *OGA* expression, rationalizing that such correlations may reflect OGT/OGA-dependent regulation of gene expression networks during development and post-natal life. Analysis of the BrainCloud transcriptomic dataset, comprising mRNA levels of over 17,000 genes across the human life span [46], revealed 4,273 genes which correlated with *OGT* (Pearson correlation coefficient > 0.5; Student’s asymptotic test *p* < 0.05; Supp. File. 1), and 4,436 genes correlated with *OGA* throughout the human lifespan (Supp. File 1). Perhaps unsurprisingly, a strong correlation was observed between *OGT* and *OGA*, consistent with previous reports highlighting the tight coupling of *OGT* and *OGA* transcription [47, 48](Fig. 1A). GO analysis of genes correlated with *OGT* identified several clusters of related biological processes, including chromatin remodeling, DNA damage response, and the stress response, all previously reported to be regulated by O-GlcNAcylation [49–51] (Fig. 1C; See Supp. File 2 for the full list of enriched GO terms). Interestingly, we observed GO terms related to the cell cycle as among the most enriched *OGT* correlated processes (26 cell cycle-related GO terms out of 155 enriched GO terms for *OGT* correlated genes), with 6% of all *OGT* correlated genes (268) annotated as being related to cell cycle progression (See Supp. File 2 for the full list of GO terms). Such genes included the M-phase regulator *CDK1* and its upstream inhibitory kinase *Wee1*, as well as the S-phase regulatory kinases *CDK4,CDK6*, and *CDK2*. *OGA* correlation analysis revealed a similar observation, with 290 cell-cycle related genes correlating with *OGA*, representing 7% of all *OGA* correlated genes across 160 cell cycle GO terms (Supp. File S1; Fig. 1C), including the tumour suppressor *RB1* and the G1/S phase repressor *CDK7*. Furthermore, there was extensive overlap between the lists of *OGT* and *OGA* correlated cell cycle genes, with 229 correlating with both O-GlcNAc cycling enzymes (Fig. 1A), perhaps expected given the dynamic and antagonistic nature of these enzymes in regulating O-GlcNAc signaling. Additionally, consistent with the overlapping nature of the *OGT* and *OGA* correlated gene lists, we observed functionally similar enriched GO terms between the *OGT* and *OGA* correlated gene sets, including genes related to double stranded break repair, histone H4 acetylation, and the stress response (Supp. File 2). To summarise, cell cycle regulators and signalling nodes are correlated with *OGT* and *OGA* throughout lifespan in the human prefrontal cortex, suggesting regulatory interplay between O-GlcNAc cycling and cellular proliferation.

**Figure 1:**
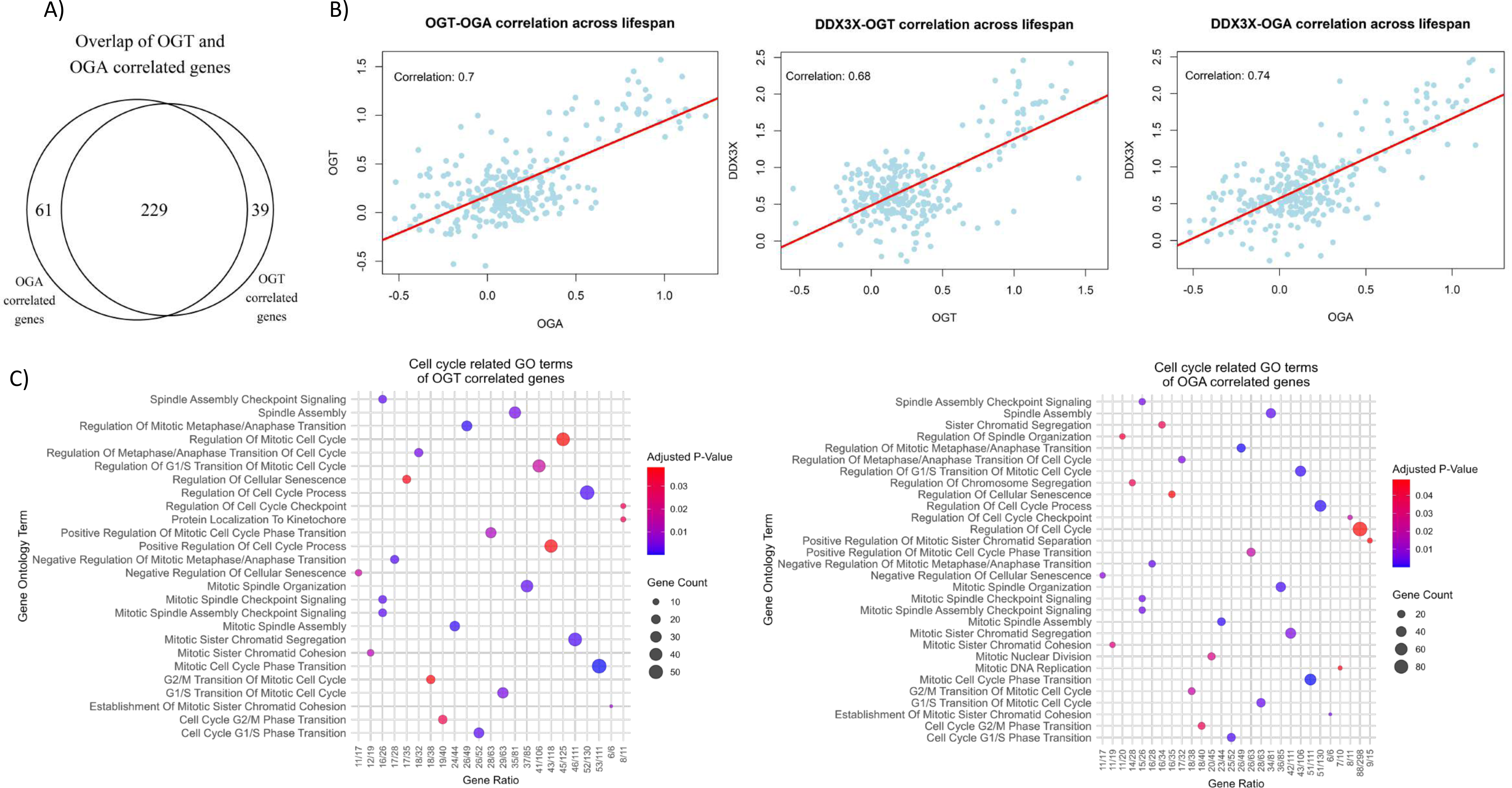
Co-expression analysis of *OGT* and *OGA* in the prefrontal cortex identifies correlations with cell cycle associated genes and processes. **(a)** Venn diagram showing the number of genes that overlap between the genes involved in the cell cycle related GO terms (adj. *P* < 0.05) and correlated with *OGT* and *OGA* (CC > 0.4 and *P*<0.05), respectively. Co-expression analysis was performed on the 17,160 genes from the BrainCloud transcriptomics dataset (n = 267) (for details see Materials and Methods). The number of cell cycle related genes correlated (CC>0.4) and anticorrelated (CC<-0.4) with *OGT* and *OGA*, respectively, are annotated (p-value<0.05). **(b)** Bubble charts showing the enriched GO BP terms related to cell cycle for both *OGT* and *OGA* correlated genes, respectively (adj. p-value<0.05). **(c)** Scatterplots showing *DDX3X*’s correlation with *OGT* (CC=0.68 and *P*<0.05) and *OGA* (CC=0.74 and *P*<0.05), respectively.

We subsequently widened our bioinformatic analysis to include genes that anti-correlated with *OGT* and *OGA* throughout life. In contrast to the *OGT* and *OGA* correlation analysis, GO terms associated with cell cycle progression were not significantly enriched, although manual inspection of *OGT* and *OGA* anti-correlated gene sets identified the gene encoding the G1/S phase-regulator *cyclin E1* (correlation coefficients of −0.74 and −0.69 respectively; Supp. File 1). We did, however, observe significant enrichment for GO terms related to synaptic transmission, in which the genes encoding SNAP25 and Vamp2, membrane tethered proteins required for pre-synaptic vesicle fusion, were observed as well as genes encoding the potassium ion channel KCNA1 and sodium/potassium ATPase ATP1A2, missense variants in both of which are associated with epilepsy and intellectual disability [52, 53]. Moreover, transcript levels of *CDK5,* a repressor of synaptic plasticity and synaptogenesis, were anti-correlated with both *OGT* and *OGA* throughout lifespan. Additionally, we observed that multiple ligand-gated ion channels essential for synaptic transmission anti-correlated with *OGT*, including *GRID1*, *GRIK4*, and *GRIN3A*, as well as *SPHN*, which is required for anchoring of mitochondria at synapses to enable synaptic transmission through localized production of ATP and Ca^2+^ buffering [54]. Future investigation will be required to determine whether these anti-correlations represent O-GlcNAc-dependent regulation of synaptic transmission and synaptogenesis through transcription of the aforementioned genes. Taken together, genes required for synaptic communication and regulation of synaptogenesis, including intellectual disability associated genes, are anti-correlated with *OGT* and *OGA*.

To investigate the possible convergence of O-GlcNAcylation and the cell cycle in intellectual disability and, by extension, OGT-CDG aetiology, we focused on the observed correlation between *OGT* and *OGA* with *DDX3X*, an RNA helicase implicated in S phase entry and intellectual disability [32, 42, 45]. We observed that *DDX3X* was among the genes most strongly correlated with *OGT* and *OGA* across lifespan in the prefrontal cortex (correlation coefficients of 0.68 and 0.74 respectively; Fig. 1B). Interestingly, DDX3X is itself an O-GlcNAc protein and missense variants in DDX3X segregate with an intellectual disability syndrome that displays clinical overlap with OGT-CDG. Although peripheral dysmorphias, namely clinodactyly and syndactyly are only observed in OGT-CDG patients, a range of endocrine, visual, cardiac, digestive, and facial morphic defects have been reported in both OGT-CDG and DDX3X ID patients (See Fig. 2 for an illustrated summary). Most notably, like OGT-CDG patients, DDX3X-ID patients often present with microcephaly, a developmental condition tightly segregating with defects in neuronal progenitor cell cycle progression [32]. Curiously, a paralogue of DDX3X residing on the Y-chromosome (DDX3Y), is able to compensate for loss of DDX3X protein and thereby prevent microcephaly in male mice, whereas female mice lack this compensatory mechanism. No such compensatory paralogue exists for OGT, which may explain the prevalence of DDX3X-ID in females, versus the high frequency of male OGT-CDG patients (Fig. 2). Aside from the apparent clinical overlap and correlated expression of *OGT* and *DDX3X*, a previous bioinformatic study proposed a link between hypo-O-GlcNAcylation of the DDX3X protein and OGT-CDG aetiology [55]. These convergent lines of evidence suggest the possibility of O-GlcNAc-dependent regulation of DDX3X, although it remains to be investigated whether any such regulation occurs at the transcriptional or post-transcriptional level. Taken together, these analyses reveal correlated expression of *OGT, OGA* and *DDX3X* in human prefrontal cortex, and highlight *DDX3X* as a candidate gene for further study.

**Figure 2:**
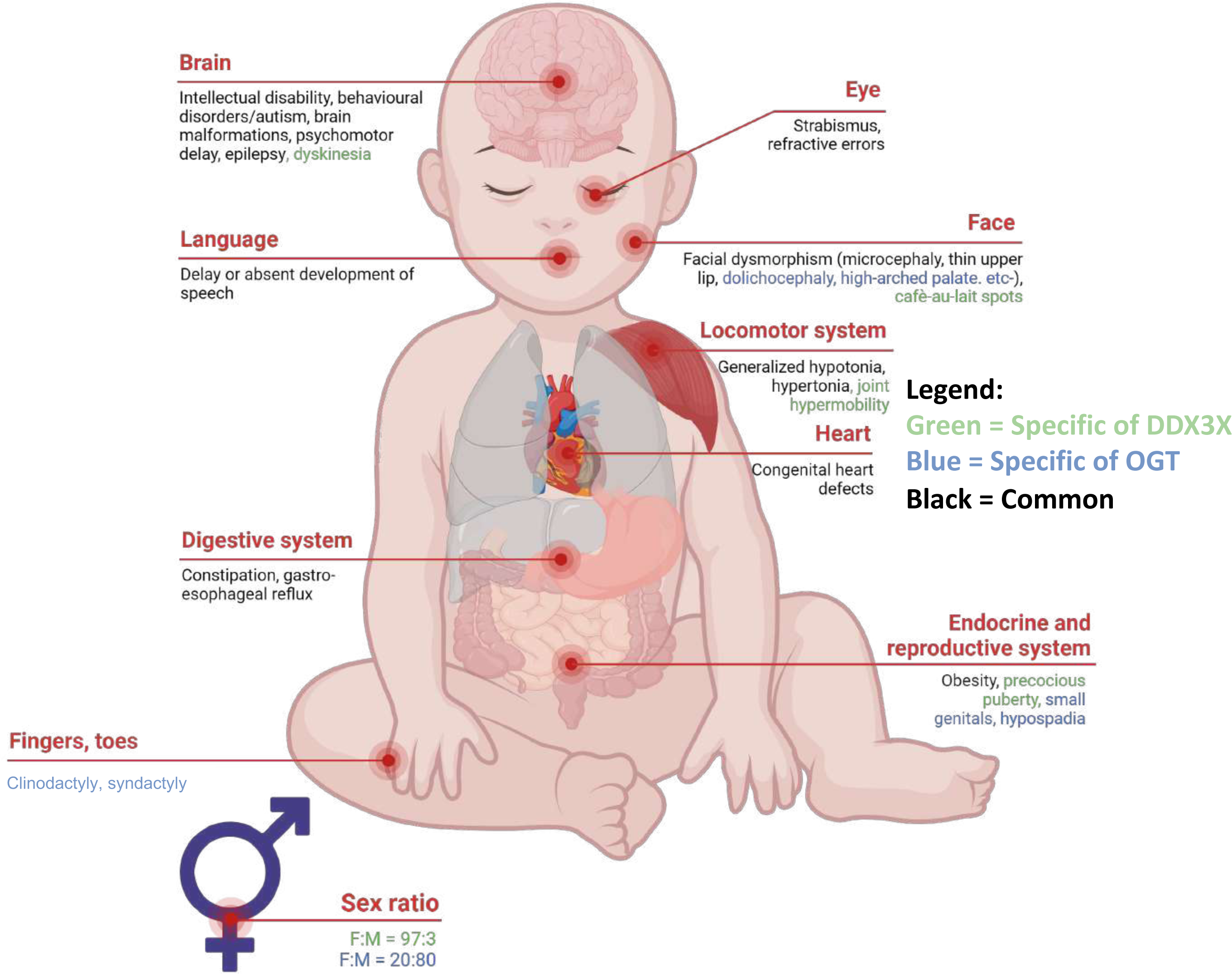
Illustrative comparison of the clinical features observed in OGT-CDG and DDX3X-XLID patients. Reported clinical features of DDX3X XLID patients were taken from [45, 83], and compared with clinical features of OGT-CDG patients in [20–25, 76]. Similarities are illustrated, along with the sex ratio for DDX3X XLID and OGT-CDG. For DDX3X XLID N = 141 patients, for OGT-CDG N = 13 patients.

### Global manipulation of O-GlcNAc homeostasis affects DDX3X levels

To investigate whether the observed correlation between *OGT* and *DDX3X* translates to the protein level, and thus whether regulatory interplay exists between these two enzymes, OGT was transiently over-expressed in HEK293T cells and DDX3X protein and mRNA levels were measured. Global elevation of O-GlcNAc levels via OGT over-expression in HEK293T cells increased endogenous DDX3X protein levels without increasing *DDX3X* mRNA levels (Fig. 3A,B), whereas over-expression of a catalytically null OGT^K852M^ mutant had no effect on DDX3X protein levels (Supp. Fig. S1). Conversely, knock-down of OGT using small interfering RNA (siRNA) resulted in reduced DDX3X protein levels (Fig. 3C). The accumulation of DDX3X protein, but not mRNA, following OGT over-expression suggested that elevated DDX3X protein levels resulted from either a post-transcriptional or post-translational effect of elevated O-GlcNAcylation on DDX3X levels. To probe this further, a metabolic labelling strategy was used to measure DDX3X O-GlcNAcylation stoichiometry in the presence or absence of exogenous OGT. Towards this end, HEK293T cells were treated with tetra-acetylated GalNAc azide (Ac_4_GalNAz). Unlike Ac_4_GlcNAz, this cell penetrant metabolic precursor is efficiently converted into UDP-GalNAz via the hexosamine salvage pathway, and subsequently epimerised to UDP-GlcNAz [56]. Importantly, the azide group is tolerated by OGT, allowing UDP-GlcNAz to be used as a donor substrate for labelling endogenous GlcNAc sites with GlcNAz. After metabolic labelling, endogenous GlcNAzylated proteins were conjugated to resolvable PEG mass tags via strain promoted azide alkyne cycloadditions (SPAAC [57]). In the absence of OGT, DDX3X shows a single resolvable band shift, compatible with mono-GlcNAcylation of the protein (Fig. 3D), whereas over-expression of OGT resulted in accumulation of poly-GlcNAcylated glycoforms of DDX3X (Fig. 3D). Taken together, these data demonstrate that manipulating O-GlcNAc homeostasis affects DDX3X protein levels, suggesting O-GlcNAc-dependent regulation of DDX3X.

**Figure 3:**
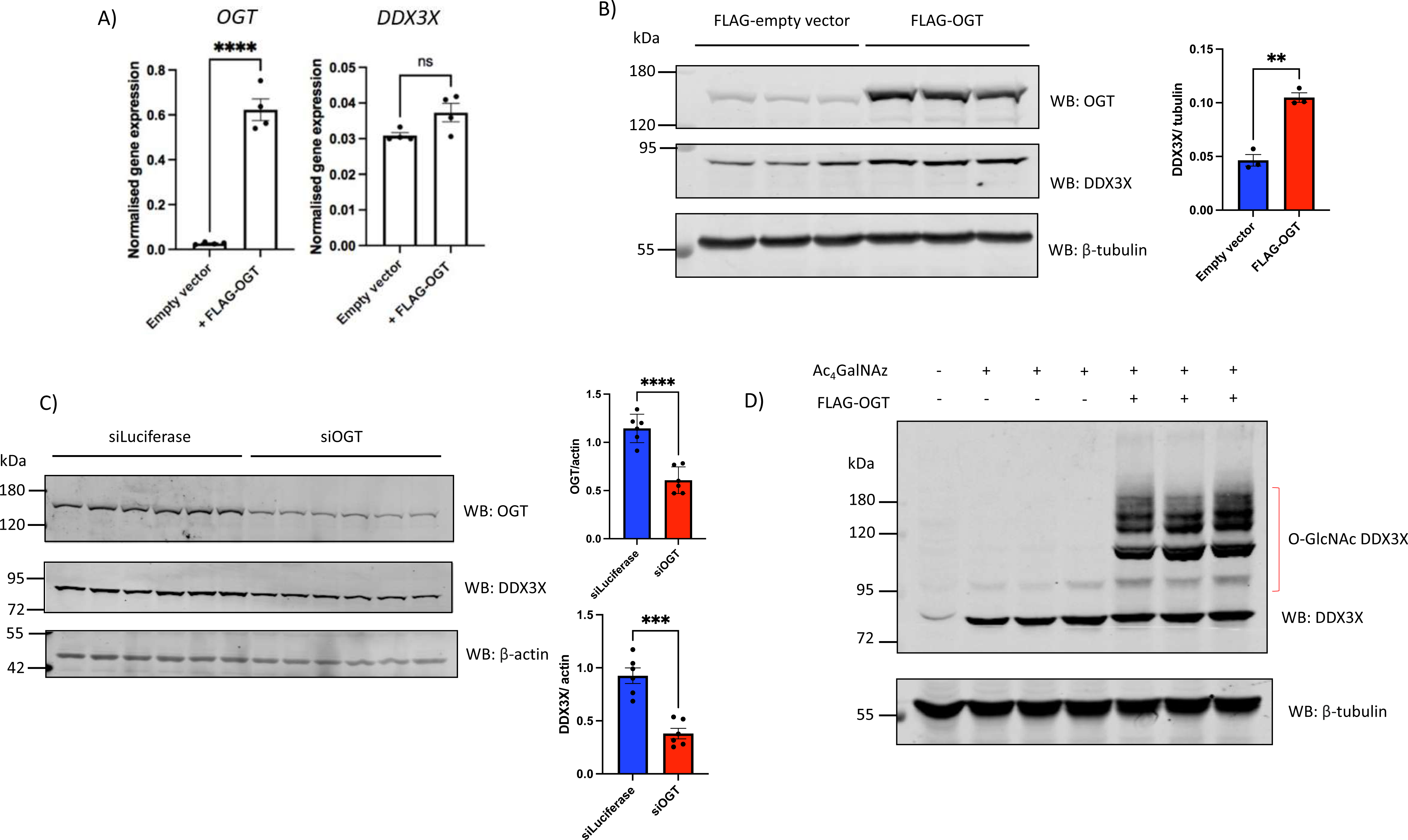
Manipulating global O-GlcNAc homeostasis elevates DDX3X protein levels and O-GlcNAcylation without altering *DDX3X* transcription. **a)** *OGT* and *DDX3X* mRNA levels after transfection of HEK293T cells with either a FLAG-empty CMV vector or equal quantity of FLAG-OGT^WT^ plasmid. *OGT* and *DDX3X* mRNA levels were normalised to the β*-actin* housekeeping gene. N = 4 technical replicates, plotted as the mean +/- S.E.M. Data were analysed using Student’s unpaired *t*-test. ns: not significant, ****: *P*<0.0001. **b)** Endogenous DDX3X protein levels after transfection of HEK293T cells in triplicate with either FLAG-empty vector or FLAG-OGT^WT^ plasmid. OGT levels are reported as a control for successful transfection with FLAG-OGT^WT^. N = 3 technical replicates. Endogenous DDX3X levels, normalised to β-tubulin, are plotted +/- S.E.M. Data were analysed using Student’s unpaired *t*-test. **: *P*<0.01. **c)** Endogenous DDX3X and OGT protein levels after transfection with siRNA targeting firefly *luciferase* (non-targeting control) or siRNA targeting human *OGT*. DDX3X and OGT levels, normalised to β-actin, are plotted +/- S.E.M. N = 6 technical replicates. Data were analysed using Student’s unpaired *t*-test. ***: *P*<0.001. **d)** HEK293T cells were transfected with either FLAG-empty vector or plasmid encoding FLAG-OGT^WT^ for 48 h. 16 h prior to cell lysis, cells were treated with 200 μM Ac_4_GalNAz to label O-GlcNAc sites with GlcNAz (see results for description of approach) or an equal volume of DMSO (vehicle only control). GlcNAz-labelled proteins were conjugated to PEG 5 kDa mass tags via strain-promoted azide alkyne cycloaddition, and reaction products analysed via DDX3X Western blot. The shifted bands within the bracket represent PEGylated DDX3X that was labelled with GlcNAz.

### Ser584 O-GlcNAcylation affects DDX3X levels

The observed accumulation of O-GlcNAcylated DDX3X following OGT over-expression suggested that O-GlcNAcylation of DDX3X results in elevated DDX3X protein levels, though the mechanism underpinning this remained unclear. To interrogate the possible site-specific effects of DDX3X O-GlcNAcylation, we mapped DDX3X O-GlcNAc sites using the metabolic labelling/ PEG mass tagging strategy described above. A previous O-GlcNAcomics study in T-lymphocytes mapped two sites of DDX3X O-GlcNAcylation at Ser584 and Ser588 in the C-terminal extension (CTE), responsible for DDX3X oligomerisation and modulation of DDX3X RNA helicase activity [58, 59]. However, the PEG mass tagging strategy used here suggests a single site of modification (Fig. 3D). To determine which of these sites makes significant contributions to DDX3X O-GlcNAcylation stoichiometry, HEK293T cells were transiently transfected with FLAG-tagged DDX3X WT, Ser584Ala, Ser588Ala, and Ser584Ala;Ser588Ala double alanine mutants, and O-GlcNAc sites were labelled with DBCO-mPEG as described above. Of the single alanine mutants analysed, FLAG-DDX3X^Ser584Ala^ showed the greatest decrease in O-GlcNAcylation stoichiometry, with the intensity of the single band shift observed for FLAG-DDX3X^WT^ reduced by 60% for FLAG-DDX3X^Ser584Ala^ (Fig. 4A, lane 4). Curiously, FLAG-DDX3X^Ser588Ala^ also showed 20% reduced O-GlcNAcylation stoichiometry (Fig. 4A, lane 5), and >90% abolition of the shifted band was only observed for the double alanine FLAG-DDX3X^Ser584Ala;Ser588Ala^ mutant (Fig. 4A, lane 6). We cannot discard the possibility that mutagenesis of Ser584 to Ala affects OGT activity on Ser588 and *vice versa*, resulting in artefactual gain or loss of OGT activity towards the remaining O-GlcNAc site. Moreover, the residual O-GlcNAcylation of FLAG-DDX3X^Ser584Ala;Ser588Ala^ suggests that additional lower occupancy O-GlcNAc sites may be present on DDX3X, an assertion supported by the 4 O-GlcNAc sites observed on endogenous DDX3X following OGT over-expression (Fig. 3D).

**Figure 4:**
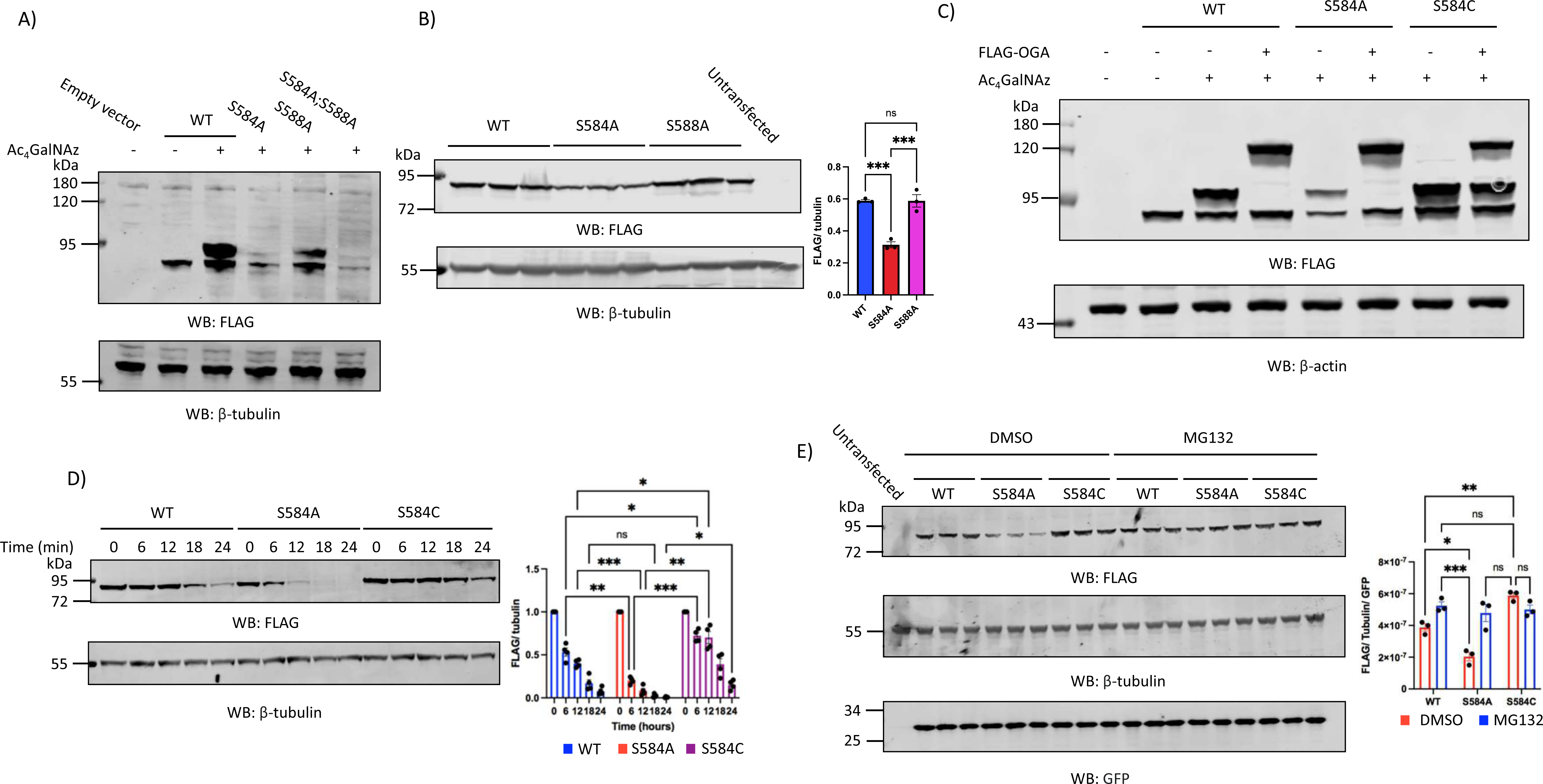
Ser584 O-GlcNAcylation protects DDX3X from degradation by the ubiquitin-proteasome system. **(a)** HEK293T cells were transfected with FLAG-empty vector, N-terminally FLAG-tagged DDX3X WT, S584A, S588A, or a S584A;S588A double alanine mutant, followed by metabolic labelling of O-GlcNAc sites with 200 μM Ac_4_GalNAz. As a negative control, a FLAG-DDX3X^WT^ transfected well was treated with an equal volume of DMSO (vehicle only control) instead of Ac_4_GalNAz. After transfection and metabolic labelling, GlcNAz-labelled proteins were conjugated to 5 kDa PEG mass tags via strain promoted azide-alkyne cycloaddition, and FLAG-DDX3X O-GlcNAcylation stoichiometry analysed by blotting for the N-terminal FLAG tag. **(b)** HEK293T cells were transfected in triplicate with FLAG-DDX3X^WT^, FLAG-DDX3X^S584A^, or FLAG-DDX3X^S588A^. Cells treated only with equivalent volumes of lipofectamine without plasmid were used as a negative control. FLAG-tagged DDX3X levels were measured via FLAG-tag Western blot and normalised to β-tubulin. Normalised FLAG-DDX3X levels are plotted as the mean +/- S.E.M. Data was analysed using one-way ANOVA (N = 3 technical replicates). ns: not significant; ***:*P*<0.001. **(c)** HEK29T cells were either transfected with one of FLAG-DDX3X^WT^, FLAG-DDX3X^S584A^, or FLAG-DDX3X^S584C^ only, or additionally co-transfected with FLAG-OGA^WT^, for 48 h. 16 h prior to cell lysis, transfected cells were treated with either 200 μM Ac_4_GalNAz or an equal volume of DMSO (vehicle only control). After co-transfection and metabolic labelling, GlcNAz-labelled sites were conjugated to 5 kDa PEG mass tags via strain promoted azide alkyne cycloaddition, and DDX3X O-GlcNAcylation stoichiometry measured via FLAG-tag Western blot. As a control for the specificity of the FLAG antibody, cells were also transfected with a FLAG-empty vector. The FLAG signal at ∼120 kDa corresponds to the expected molecular weight of transfected FLAG-OGA^WT^. **(d)** HEK293T cells were transfected with either FLAG-DDX3X^WT^, FLAG-DDX3X^S584A^, or FLAG-DDX3X^S584C^ prior to addition of 100 μg/mL cycloheximide for 0,6,12,18 or 24 h. The turnover of each FLAG-DDX3X construct was measured by normalising the FLAG signal at each time-point to both β-tubulin and the FLAG signal at T = 0. Data are plotted as the mean +/- S.E.M. Data was analysed using two-way ANOVA (N = 3 technical replicates). ns: not significant, *: *P* < 0.05, **: *P* < 0.01, ***: *P* < 0.001. **(e)** HEK293T cells were transfected with either a FLAG-empty vector or a dual expression vector encoding FLAG-DDX3X^WT^, FLAG-DDX3X^S584A^, or FLAG-DDX3X^S584C^, and a 3’ P2A sequence that is co-translationally cleaved followed by the open reading frame for GFP (see results for a description of the P2A-GFP vector). Transfected cells were treated with either 10 μM MG132 or an equal volume of DMSO for 16 h prior to cell lysis. Transfected DDX3X levels were quantified via normalisation of the FLAG-tag signal to both the β-tubulin housekeeping protein and the co-expressed GFP reporter. Normalised FLAG-DDX3X levels are plotted as the mean +/- S.E.M. Data were analysed using a one-way ANOVA (N = 3 technical replicates). ns: not significant; *:*P*<0.05; \*\**P*<0.01; \*\*\**P*<0.001.

In addition to providing information on O-GlcNAc site localisation and stoichiometry, the single and double alanine mutants also suggested an effect on the steady state levels of DDX3X in cells, as the Ser584Ala single and Ser584Ala;Ser588Ala double alanine mutants both expressed at lower levels than WT (Fig. 4A). Indeed, repetition of transfection experiments in triplicate showed a 60% reduction in steady state expression of FLAG-DDX3X^Ser584Ala^ and FLAG-DDX3X^Ser584Ala;Ser588Ala^, but not FLAG-DDX3X^Ser588Ala^ (Fig. 4B). This 60% decrease in FLAG-DDX3X^Ser584Ala^ protein level was proportional to the decrease in DDX3X O-GlcNAcylation stoichiometry after alanine mutagenesis of Ser584 (Fig. 4A). Melting curves of DDX3X^WT^ and DDX3X^Ser584Ala^ recombinant protein, expressed in *E. coli*, did not differ (Supp. Fig. S2), indicating that the differences in FLAG-DDX3X^WT^ and FLAG-DDX3X^Ser584Ala^ expression levels did not stem from the alanine mutation disrupting the protein fold. As an additional control, transfection of HEK293T cells with FLAG-DDX3X plasmids where the 3’ end of the *DDX3X* ORF was fused via a co-translationally cleaved P2A linker to GFP, revealed no differences in GFP protein levels (Fig. 4E), suggesting that the observed reduction in steady state FLAG-DDX3X^Ser584Ala^ levels did not stem from differential transfection or translation efficiencies and therefore results from loss of O-GlcNAcylation. To summarise, Ser584 O-GlcNAcylation affects DDX3X protein levels.

### Ser584 O-GlcNAcylation prevents DDX3X degradation by the ubiquitin-proteasome system

The effects of protein O-GlcNAcylation on proteostasis have been well documented, with examples of O-GlcNAcylation inhibiting protein aggregation, increasing protein thermostability, and inhibiting degradation via the ubiquitin-proteasome system (UPS), reported previously [60–64]. The observation that over-expression of OGT elevated DDX3X O-GlcNAcylation and protein levels without affecting *DDX3X* transcription, paired with observed reductions in the levels of the Ser584Ala mutant, suggested a post-translational mechanism through which O-GlcNAc stabilises DDX3X. To investigate this hypothesis, FLAG-DDX3X^WT^ and FLAG-DDX3X^Ser584Ala^ were transiently over-expressed, and their turnover rates *in cellulo* analysed after blocking translation with the protein synthesis inhibitor cycloheximide. The rate of FLAG-DDX3X^Ser584Ala^ turnover was 3-fold higher than for FLAG-DDX3X^WT^ (Fig. 4D), with no detectable FLAG-DDX3X^Ser584Ala^ after 18 h cycloheximide treatment, in contrast to FLAG-DDX3X^WT^ where 20% of the initial pool of FLAG-DDX3X^WT^ could be detected at the same time point (Fig. 4D). Moreover, treatment of FLAG-DDX3X transfected cells with the proteasome inhibitor MG132 rescued FLAG-DDX3X^Ser584Ala^ expression (Fig. 4E), suggesting the increased turnover of FLAG-DDX3X^Ser584Ala^ stemmed from increased degradation via the UPS. As an additional control, we installed an OGA-resistant S-GlcNAc analogue of O-GlcNAc at Ser584 via mutation of this residue to Cys. This approach exploits a recently discovered OGT promiscuity towards Cys, combined with the observation that S-GlcNAcylated Cys are resistant to OGA hydrolysis, thus resulting in site-specific elevation/installation of GlcNAcylation in the context of an otherwise unaltered O-GlcNAcome [65–67]. FLAG-DDX3X^S584C^ demonstrated a 30% increase in O-GlcNAcylation stoichiometry relative to FLAG-DDX3X^WT^, and whilst co-transfection with FLAG-OGA^WT^ abolished O-GlcNAcylation of FLAG-DDX3X^WT^ and FLAG-DDX3X^S584A^, the intensity of the shifted band was only reduced by 20% for FLAG-DDX3X^S584C^ (Fig. 4C). The 20% reduction in FLAG-DDX3X^S584C^ O-GlcNAcylation stoichiometry after OGA over-expression could reflect removal of Ser588 O-GlcNAcylation with preservation of S-GlcNAc at Cys584. Hyper-S-GlcNAcylated FLAG-DDX3X^Ser584Cys^ showed higher steady state expression levels than FLAG-DDX3X^WT^ (Fig. 4E), which could not be elevated further by the addition of MG132. Hyper-S-GlcNAcylated FLAG-DDX3X^Ser584Cys^ also showed a reduced turnover rate in the cycloheximide assay compared to FLAG-DDX3X^WT^ (Fig. 4D). Take together, these data suggest that Ser584 O-GlcNAcylation protects DDX3X from degradation by the UPS.

### Ser584 O-GlcNAcylation slows DDX3X aggregation kinetics in vitro

We next investigated how Ser584 O-GlcNAcylation might prevent DDX3X turnover *in cellulo*. Given that Ser584 is within the intrinsically disordered C-terminal extension (CTE) of DDX3X, it is possible that Ser584 O-GlcNAcylation prevents aggregation of DDX3X and its subsequent targeting for degradation by the UPS. To investigate this possibility, we immunoprecipitated transiently transfected FLAG-DDX3X^WT^ and FLAG-DDX3X^S584A^ from HEK293T lysates. Immunoprecipitates were subsequently incubated at 37 °C to promote aggregation of DDX3X. At each time-point investigated, we observed an approximately 2-fold higher abundance of FLAG-DDX3X^WT^ in the soluble fraction compared to hypo-O-GlcNAcylated FLAG-DDX3X^S584A^ and, conversely, faster accumulation of GlcNAc-deficient FLAG-DDX3X^S584A^ in the insoluble fraction compared to O-GlcNAcylated FLAG-DDX3X^WT^ (Supp. Fig. S3, compare to spiked pre-immune rabbit IgG and lysozyme loading controls). Given the faster depletion of soluble unmodified FLAG-DDX3X^S584A^ compared to O-GlcNAcylated FLAG-DDX3X^WT^, paired with our previous observation that alanine mutagenesis at position 584 does not alter DDX3X protein folding (Supp. Fig. S2), these data suggest that DDX3X Ser584 O-GlcNAcylation may prevent aggregation and reduce DDX3X targeting to the UPS.

### Loss of DDX3X Ser584 O-GlcNAcylation impairs cellular proliferation through stalling of the G1/S phase transition

DDX3X is a multi-functional RNA helicase, with RNA helicase-dependent and independent roles in cellular proliferation, the stress response, and neurite outgrowth [68, 69]. A previous study in immortalised cell lines linked DDX3X to cell cycle progression through the ability of DDX3X to promote *cyclin E1* translation, thus permitting entry into S phase of the cell cycle. Moreover, as discussed previously, DDX3X loss of function causes microcephaly partially through stalled cell cycle progression of neuronal progenitors, marking cell cycle progression as a process that is dysregulated in DDX3X ID. We therefore next investigated whether loss of Ser584 O-GlcNAcylation increases DDX3X turnover sufficiently to impair cell cycle progression. We initially investigated whether loss of Ser584 O-GlcNAcylation noticeably impaired cellular proliferation through an siRNA-rescue strategy. Endogenous *DDX3X* was knocked-down, and knock-down (KD) cells were co-transfected with either siRNA resistant (from here-on, SIR) FLAG-DDX3X^WT^ or FLAG-DDX3X^Ser584Ala^. Transfection of these constructs was titrated to rescue total DDX3X levels to those observed in cells transfected with siRNA targeting *firefly luciferase* (non-targeting control) to prevent artefactual results due to supra-physiological levels of DDX3X. After 72 h of transfection, *DDX3X* KD cells showed reduced live cell counts of 79% and 84% by trypan blue staining and MTT assays relative to the non-targeting control, respectively (Fig. 5C, Supp. Fig. S6). Co-transfection with SIR FLAG-DDX3X^WT^ rescued this reduction in viability, whereas SIR FLAG-DDX3X^Ser584Ala^ did not (Fig. 5C, Supp. Fig S4). Western blotting of cell lysates revealed the same effect of loss of Ser584 O-GlcNAcylation as observed in previous experiments (Fig. 5A), with KD reducing DDX3X protein levels by 61%, and SIR FLAG-DDX3X^WT^ but not SIR FLAG-DDX3X^Ser584Ala^ rescuing total DDX3X levels (Fig. 5A). Levels of cyclin E1, the translation of which requires DDX3X activity, were reduced by 71% following *DDX3X* KD, and this was rescued by SIR FLAG-DDX3X^WT^ but not SIR FLAG-DDX3X^Ser584Ala^, suggesting an S-phase entry defect upon loss of DDX3X Ser584 O-GlcNAcylation (Fig. 5A). These data collectively pointed suggest possible defects in cell cycle progression, which we next investigated by flow cytometry. *DDX3X* KD resulted in a 63% increase in the G1 population, and concomitant reductions in the percent of cells in S and G2/M phase (by 29% and 67% respectively, Fig. 5B). Co-transfection with SIR FLAG-DDX3X^WT^ restored the cell cycle distribution to that observed in the non-targeting control (Fig. 5B). However, SIR FLAG-DDX3X^Ser584Ala^ cells showed accumulation of the G1 population similar to that observed in *DDX3X* KD cells (Fig. 5B). Indeed, a 65% increase in the G1 population was observed, alongside reduced cell cycle occupancy in S and G2/M phases, suggesting turnover of GlcNAc-deficient DDX3X^S584A^ was sufficiently high to impair S phase entry. Taken together these findings suggest that loss of Ser584 O-GlcNAcylation accelerates UPS-dependent turnover of DDX3X to such an extent that progression from G1 into S phase of the cell cycle is impaired, resulting in reduced cellular proliferation.

**Figure 5:**
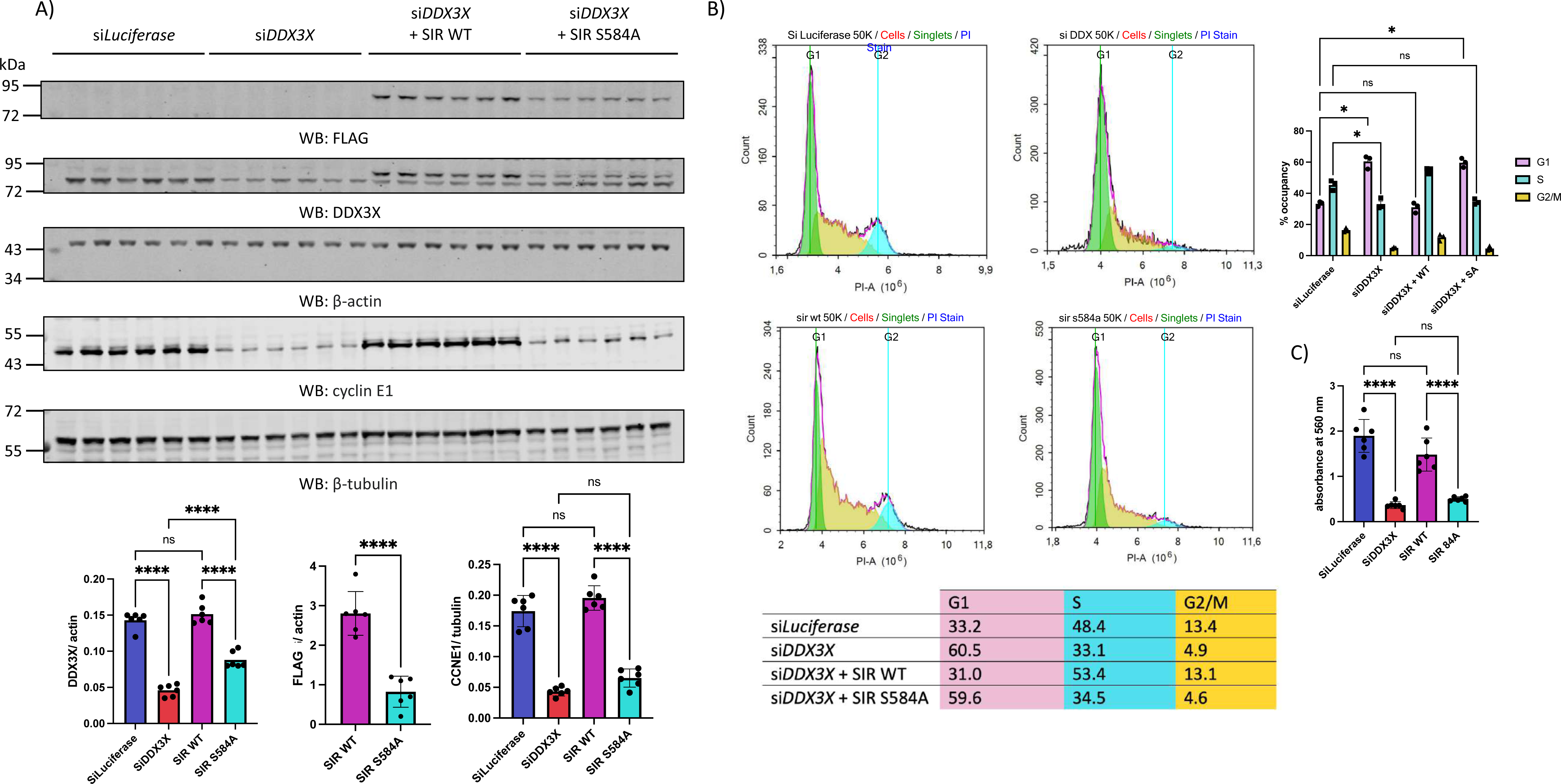
Loss of DDX3X Ser584 O-GlcNAcylation impairs S-phase entry and stalls cellular proliferation. **a)** HEK293T cells were transfected with siRNA against either firefly *luciferase* (non-targeting control) or endogenous human *DDX3X*. In parallel, *DDX3X* knock-down cells were also co-transfected with either siRNA-resistant (SIR) FLAG-DDX3X^WT^ or SIR FLAG-DDX3X^S584A^. After 72 h, live cell numbers were counted (see Supp. Fig. S6), and cells lysed for analysis of FLAG-DDX3X, total (endogenous and FLAG-DDX3X) DDX3X, and cyclin E1 levels by Western blotting. The FLAG, total DDX3X, and cyclin E1 signals, normalised to either β-actin (FLAG, total DDX3X) or β-tubulin (cyclin E1), are plotted as the mean +/- S.E.M. Normalised total DDX3X and cyclin E1 data were analysed by one-way ANOVA (N = 6 technical replicates). ns: not significant; ****: *P* < 0.0001. Normalised FLAG-DDX3X data were analysed using Student’s unpaired *t*-test (N = 6 technical replicates). ****: *P* < 0.0001. Data presented are from one of two independent experiments. **b)** Cell cycle distribution of HEK293T cells transfected with si*luciferase* or si*DDX3X*, or co-transfected with si*DDX3X* and either SIR FLAG-DDX3X^WT^ or FLAG-DDX3X^S584A^ as determined by flow cytometry. Cells in G1, S, and G2/M are highlighted in green, yellow, and cyan respectively. The average occupancy of each cell population in G1,S and G2/M is listed in table format. Data from 3 independent experiments was analysed using two-way ANOVA. ns: not significant, \**P*<0.05. **c)** 1 x 10^5^ cells were transfected with siRNA against *luciferase* or *DDX3X*, with si*DDX3X* cells additionally co-transfected with SIR FLAG-DDX3X^WT^ or SIR FLAG-DDX3X^S584A^. After 72 h, media was aspirated and replaced with 1 mL of 0.5 mg/mL thiazolyl blue tetrazolium bromide dissolved in cell media for 4 h. After 4 h, DMSO was added to solubilise formazan crystals, and the absorbance at 560 nm measured. Absorbance measurements are plotted as the mean +/- S.E.M. N = 6 technical replicates. Data were analysed using one-way ANOVA. ns: not significant, ****:*P*<0.0001.

### DDX3X proteins levels are reduced in a mouse model of OGT-CDG

Missense variants in OGT segregate with a novel X-linked intellectual disability syndrome, the O-GlcNAc transferase congenital disorder of glycosylation (OGT-CDG), a clinically heterogenous syndrome that remains to be mechanistically dissected [27]. Previously, we identified a group of neuronal, ID-associated, O-GlcNAc proteins that could contribute to distinct aspects of OGT-CDG, which included DDX3X [55], and we identified correlations between expression of *OGT*, *DDX3X* and its target *cyclin E1* here (Fig. 1C; Supp. File 1). Given that we have demonstrated that O-GlcNAcylation inhibits DDX3X turnover by the UPS, and that loss of this modification is sufficient to impair at least one DDX3X-regulated cellular process, we sought to investigate whether DDX3X protein levels may be reduced in mouse models of OGT-CDG, where global O-GlcNAc levels are reduced in brain [70]. *DDX3X* mRNA levels were unaltered in brain lysates derived from these mice (Fig. 6B). Interestingly, in the brains of OGT^N648Y^ mice DDX3X protein were reduced by 50% (Fig. 6A). Given the previously reported requirement for DDX3X in *cyclin E1* translation, and the documented role of cyclin E1 in learning and memory, we also probed cyclin E1 protein levels and similarly observed a 66% decrease in OGT^N648Y^ adult mouse brains. As reductions in DDX3X protein levels were only observed in OGT^N648Y^ mice, DDX3X was immunoprecipitated from OGT^N648Y^ mouse brain lysates and blotted with a pan-specific O-GlcNAc antibody (RL2). Despite the observed reduction in DDX3X protein levels, O-GlcNAcylation was not reduced on immunoprecipitated DDX3X by RL2 Western blotting (Fig. 6D). It is possible that the non-O-GlcNAcylated fraction of DDX3X is subject to rapid turnover by the UPS, resulting in more of the O-GlcNAcylated fraction remaining, which could explain the equal levels of RL2 immunoreactivity on immunoprecipitated DDX3X from wild-type and OGT-CDG mice. Additionally, it is important to note that RL2 was initially raised against proteins of the nuclear pore complex and was only later found to bind O-GlcNAc [71, 72]. Therefore, it is not clear whether the RL2 signals observed are *bona fide* O-GlcNAc signals or the result of RL2 binding to the protein backbone. To summarise, levels of DDX3X and its target cyclin E1 are reduced in OGT-CDG mouse brains, pointing to dysregulated DDX3X proteostasis in OGT-CDG.

**Figure 6:**
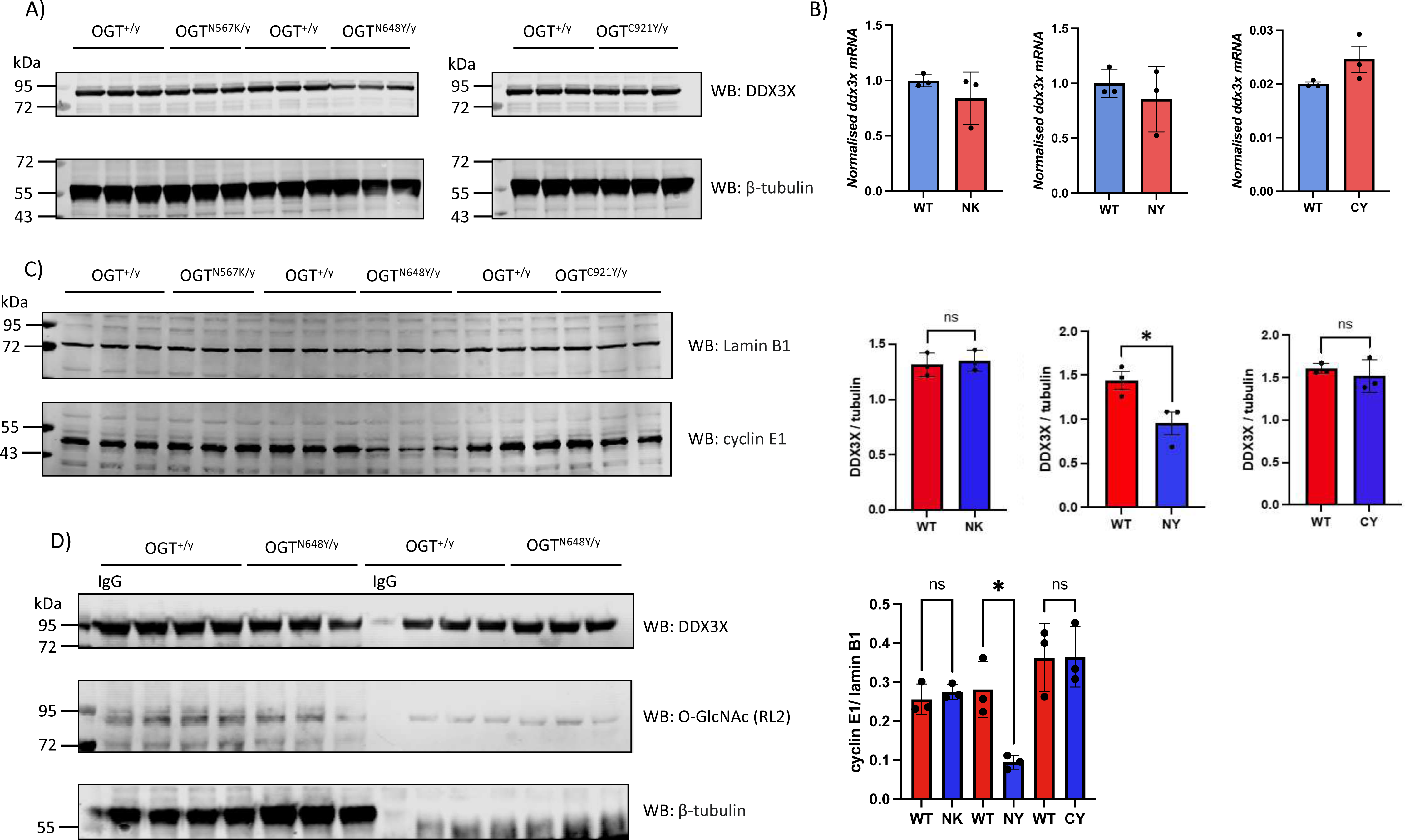
DDX3X expression, but not O-GlcNAcylation stoichiometry, is reduced in the brains of OGT^N648Y^ mice. **a)** Endogenous DDX3X protein levels were probed via Western blotting of whole brain homogenates from adult male wild type or hemizygous OGT-CDG mouse lines. DDX3X signals were normalised to β-tubulin. Data were analysed using Student’s unpaired T-test (N = 3 biological replicates). ns: not significant. *: *P*<0.05. **b)** *DDX3X* mRNA levels from whole brain homogenates of adult male wild type or hemizygous OGT-CDG mouse lines were quantified via RT-qPCR and normalised to the average Cq value of the housekeeping genes *18S*, *actin* and *PGK.* Normalised gene expression is plotted as the mean +/- 1 SD. Data were analysed using Student’s unpaired *t*-test (N = 3 biological replicates). ns: not significant. **c)** Cyclin E1 protein levels, normalised to lamin B, from whole brain homogenates of adult male wildtype and OGT-CDG mouse lines. Normalised cyclin E1 protein levels are plotted as the mean +/- S.E.M. Data were analysed using one-way ANOVA (N = 3 biological replicates). ns: not significant, *:*P*<0.05. **d)** Immunoprecipitation of DDX3X from whole brain lysates of OGT^+/y^ and OGT^N648Y/y^ mice. O-GlcNAc levels on immunoprecipitated DDX3X were probed using the pan-specific O-GlcNAc antibody RL2.

## Concluding remarks

Over 5000 nuclear, cytoplasmic, and mitochondrial proteins are modified with O-GlcNAc [2], a functionally diverse PTM that has been reported to modulate a plethora of cellular and developmental processes [14, 17, 64, 73–75]. A comprehensive understanding of the role O-GlcNAc plays in human physiology is impaired by the lack of information on the site-specific effects of this modification on OGT substrates and the consequent effects on cellular processes. Moreover, mutations in OGT give rise to a clinically heterogenous intellectual disability syndrome – OGT-CDG – thought to involve altered O-GlcNAc levels on key substrates that play a role in neurodevelopment/neurophysiology [20–25, 76]. The widespread distribution of O-GlcNAc across over 5000 proteins and diverse cellular processes complicates prediction of which cellular processes are pathologically altered in OGT-CDG. As a starting point towards understanding which cellular and developmental processes are pathologically disrupted in OGT-CDG, we chose to focus on identifying candidate conveyors of the microcephaly observed in OGT-CDG [69, 77]. The development of microcephaly is closely linked with dysregulated cell cycle progression of neuronal progenitors during corticogenesis [32, 43], indicating that cell cycle progression is affected in OGT-CDG patients during this critical developmental window. How O-GlcNAcylation regulates cell cycle progression, however, is poorly understood, particularly in the earlier stages of the cell cycle prior to mitosis. Thus, rational prediction of candidate proteins, the hypo-O-GlcNAcylation of which may drive microcephaly in OGT-CDG, is challenging. Indeed, to date all symptoms described in OGT-CDG patients and phenotypes in model organisms of the disease remain unexplained at the molecular level, with no dysregulated signalling pathways described to date.

As a starting point towards identifying possible molecular pathways disrupted in OGT-CDG patients, we analysed transcriptomic data from the human prefrontal cortex to identify genes and biological processes that correlated strongly with the O-GlcNAc cycling enzymes, rationalising that such correlations may reflect regulatory interplay between O-GlcNAc and key cellular and neurobiological processes. Gratuitously, we observed that expression of the master M phase regulator *CDK1,* and its upstream regulatory phosphatase *CDC25*, were correlated with *OGT* in the human prefrontal cortex. CDC25 activates CDK1 by dephosphorylation of Ser14 and Tyr15 on CDK1 to promote M phase progression [78], and the observation that both *CDK1* and *CDC25* positively correlate with *OGT* expression suggests a layer of O-GlcNAc-dependent transcriptional regulation in promoting M phase progression. Indeed, a previous study reported delayed M-phase progression following OGA over-expression [44, 45]. Future work focused on investigating the putative regulatory effects of O-GlcNAcylation on *CDK1* transcription and its regulation by CDC25 may shed light on the poorly understood role of O-GlcNAc in M-phase progression. Aside from M phase, we observed positive correlations between the master G1/S-phase regulators *CDK4*, *CDK6,* and *CDK2* with *OGT*. The CDK4/6-cyclin D and CDK2-cyclin E complexes function maximally in early and late G1 phase respectively and are required for hyperphosphorylation of retinoblastoma (Rb) [79], amongst other targets [80]. Rb hyperphosphorylation liberates the transcription factor E2F from a repressive Rb-E2F complex enabling transcription of genes required for DNA replication [80]. Previous work has pointed to a convergence between O-GlcNAc and the G1/S transition of the cell cycle, with OGA over-expression reducing Rb phosphorylation and uncoupling cyclin E1 expression from G1 phase progression [32]. Additionally, global depression of O-GlcNAc through OGT inhibition and OGA over-expression both result in delayed S-phase entry [40]. Despite these studies suggesting an important role for O-GlcNAc in S-phase entry, how individual O-GlcNAcylation events regulate S-phase entry is poorly understood, especially at the level of site-specific effects.

The transcriptomic analysis described here identified the cell cycle regulator and ID protein *DDX3X* as one of the strongest *OGT* correlated genes in the prefrontal cortex. DDX3X is an RNA helicase of the DEAD box family that regulates cell cycle progression at both the translational and transcriptional level and is critical for translation of *cyclin E1* and co-regulating transcription of the CDK inhibitor *p21* [42, 81, 82]. Indeed, loss of DDX3X results in either G1/S phase stalling or global delays in cell cycle progression depending on the cell type studied [32, 42, 43]. Importantly, missense, nonsense, and frameshift variants in DDX3X result in an ID-syndrome that, similarly to OGT-CDG, frequently presents with microcephaly [44, 45, 83]. The microcephaly in DDX3X-ID likely stems from impaired neuronal progenitor proliferation during corticogenesis, as demonstrated by elegant DDX3X loss-of-function experiments in mice [32]. Given the correlation between *DDX3X* and *OGT*, and our interest in identifying novel cell cycle regulators, the hypo-O-GlcNAcylation of which may contribute to microcephaly, we focused on investigating the possible regulatory interplay between O-GlcNAc and DDX3X. Our subsequent work identified Ser584 O-GlcNAcylation as a proteostatic regulator of DDX3X activity, with loss of Ser584 O-GlcNAcylation inhibiting turnover of DDX3X by the ubiquitin-proteasome system. Importantly, the increased turnover rate of GlcNAc-deficient DDX3X^Ser584Ala^ was sufficient to impair S-phase entry. Indeed, the observed accumulation of DDX3X^Ser584Ala^ cells in G1 phase, but not S-phase, paired with the reduced population of G2/M phase cells points to delayed transitioning through the G1/S checkpoint following loss of DDX3X Ser584 O-GlcNAcylation, as opposed to delayed release of cells into G1 following mitosis. Supporting this is the observation that cyclin E1 levels are downregulated in DDX3X^Ser584Ala^ cells, consistent with the previously documented role of DDX3X in promoting *cyclin E1* mRNA translation [42]. Given the essential role cyclin E1 plays in CDK2 activation, DDX3X O-GlcNAcylation may contribute to CDK2 activation and downstream gene expression networks in late G1, thus explaining the S-phase entry defect. Previously, chemical inhibition of OGT using acetyl-5S-GlcNAc in MCF-7 breast cancer cells was reported to cause a reduction in the percentage of S-phase cells without affecting DNA replication rate, indicative of impaired G1/S phase transitioning as opposed to prolonged S-phase duration [40], although the authors were unable to determine which O-GlcNAc signalling nodes were dysregulated to convey this effect. Our data here indicate that DDX3X may well be one the conveyors of the stalled G1/S transition observed in this previous study. It is, however, likely that OGT inhibition additionally affects S-phase entry via other mechanisms unrelated to DDX3X, with DDX3X acting as one of many O-GlcNAc-dependent regulatory mechanisms that have yet to be identified. Relevant to this point is the prior observation that global O-GlcNAc levels decrease as cells enter S-phase [84], whereas O-GlcNAcylation of DDX3X is required for S-phase entry, indicating O-GlcNAc does not solely act as a promoter or repressor of S-phase entry. Indeed, other DEAD box RNA helicases, replication licensing factors, and cytoskeletal proteins are dynamically O-GlcNAcylated in late G1, although how O-GlcNAc modulates the activity of these proteins to regulate the G1/S transition is unclear [84]. To our knowledge, this study reports the first identification of a single O-GlcNAcylation event that regulates the G1/S transition, emphasising the paucity of information on how O-GlcNAc regulates this critical point of the cell cycle and the need to further study this poorly understood area of biology.

Having characterised the proteostatic and cell cycling effects of DDX3X Ser584 O-GlcNAcylation *in cellulo*, we investigated whether DDX3X protein levels were reduced in mouse models of catalytically deficient OGT-CDG variants, hypothesising that DDX3X hypo-O-GlcNAcylation could result in increased DDX3X turnover. Although we did not observe reduced DDX3X O-GlcNAcylation by RL2 Western blotting, we observed a substantial decrease in DDX3X protein without gross effects on DDX3X transcription in the brains of OGT^N648Y^ mice. The inability to detect DDX3X hypo-O-GlcNAcylation may reflect depletion of non-O-GlcNAcylated DDX3X by the proteasome, or cell-type and tissue-specific differences in how O-GlcNAc regulates protein activity. Indeed, DDX3X turnover is regulated by K48-linked ubiquitylation catalysed by the RNF39 and CUL3 E3 ligases [85, 86], and it is possible that the activity of these and possibly other E3 ligases and kinases is elevated in OGT-CDG mice, resulting in loss of DDX3X protein. It is also presently unclear as to why DDX3X is only reduced in the N648Y brain, and not in the C921Y or N567K mouse brains. A possible explanation for this lay in the different manners by which each OGT-CDG variant affects catalytic activity. For example, whereas the C921Y variant is predicted to disrupt binding of the UDP-GlcNAc donor substrate [25], the N567K variant disrupts acceptor substrate binding through partial steric blockade of the −2 subsite when bulky side chains are present [23]. Thus, catalytically deficient OGT-CDG variants may result in hypo-O-GlcNAcylation and dysregulation of different signalling cascades and regulatory networks, which would explain our observations with DDX3X here. Future proteomic and glycoproteomic analyses of OGT-CDG mouse models will be required to investigate this assertion, and thereby identify variant-specific as well as commonly dysregulated signalling nodes.

As discussed previously, DDX3X missense, frameshift, and nonsense variants cause an ID syndrome that presents with autistic spectrum behaviours, delayed language acquisition, eye malformation, and microcephaly [44, 45, 83]. The reduced levels of DDX3X protein in the brains of OGT^N648Y^ mice, which display microcephaly, suggest that loss of DDX3X activity may contribute to the microcephaly observed in these mice as well as OGT-CDG patients [27]. Indeed, we detected a 50% decrease in DDX3X protein levels in OGT^N648Y^ mice, and loss of 70% of DDX3X protein caused microcephaly in female DDX3X loss-of-function mice through impaired cell cycling of neuronal progenitors during embryonic corticogenesis [32]. However, whether DDX3X protein levels are reduced in the radial glial cells and intermediate progenitors, which give rise to mature neurons during corticogenesis, was not investigated in this study, with the DDX3X levels being quantified in the brains of adult mice. Investigating DDX3X proteostasis during corticogenesis in OGT-CDG mice, and determining whether neuronal progenitor cell cycling is impaired, could provide explanations at both the cellular and molecular level for the occurrence of microcephaly in OGT-CDG. It is also noteworthy that we observe reductions in cyclin E1 levels in OGT-CDG mouse brains. Aside from its established role in cell cycle progression, cyclin E1 regulates synapse function and number by sequestering CDK5 into an inactive complex [87]. Conditional knock-out of *cyclin E1* in mice consequently results in impaired learning and memory due to unconstrained CDK5 activity [87]. Although it is unclear whether the observed reductions in cyclin E1 are a direct consequence of reduced DDX3X activity, it is noteworthy that cyclin E1 and DDX3X are both reduced exclusively in the brains of OGT^N648Y^ mice. The dual observations that the microcephaly associated protein DDX3X and synaptic regulator cyclin E1 are both reduced in OGT-CDG mice suggests a combination of neurodevelopmental and neurophysiological contributions to the aetiology of this disease, and will drive future research and identification of therapeutic strategies for this devastating disease.

## Materials & Methods

### Analysis of prefrontal cortex RNAseq data

To investigate temporal *OGT* and *OGA* co-expression in the human prefrontal cortex as a strategy to identify candidate genes involved in the same neurobiological pathways, the BrainCloud dataset containing human postmortem gene expression data from brain tissue was used [46]. The BrainCloud dataset was produced from transcriptomic analysis of RNA isolated from the dorsolateral prefrontal cortex Brodmann areas 46/9 grey matter tissue from foetal ages through ageing in 267 individuals with no severe neuropathological, neurological or neuropsychiatric diagnoses, across the human lifespan. Transcriptomic data were downloaded from the GEO repository (GSE30272) together with the annotation file and the demographic data. The different probes targeting the same gene were aggregated to their mean value, resulting in 17,160 genes being included in the subsequent analyses. All analyses were performed using RStudio (R v4.3.0; R Core Team 2023). The Pearson correlation coefficient (PCC) was calculated for *OGT* and *OGA* and all other genes, respectively, using the *cor()*-function from the base R-package stats (v3.6.2). Afterwards, gene lists of correlation and anti-correlation were extracted, including only genes with a PCC>=0.4 (or PCC<=-0.4) and *P* < 0.05, Student’s asymptotic test. To identify which of the genes were specifically involved in cell cycle-related processes, the gene lists were submitted to the Enrichr online resource to perform gene ontology enrichment analysis for biological processes [88]. Enriched terms were only considered significant if *P* < 0.05 after Benjamini-Hochberg correction for multiple comparisons.

### Cloning of DDX3X and OGT constructs for HEK293T transfection and E. coli expression

A PCR product containing the Human ORF for DDX3X was obtained directly from RNA using the Primescript High Fidelity RT-PCR kit from Takara. The PCR primers introduced *Bam*HI and *Not*I sites at the 5’ and 3’ ends respectively. The fragment was cloned into pCMV-FLAG as a *Bam*HI-*Not*I restriction fragment. The insert was confirmed by DNA sequencing. Site directed mutations at S584A, S584C, S588A and the double mutation S584A;S588A were introduced using site directed mutagenesis based on the Quickchange mutagenesis kit, but KOD polymerase was used instead of Pfu. The sequence was verified again by sequencing.

To confer siRNA immunity on the DDX3X construct a geneblock was designed (from IDT – Integrated DNA Technologies) containing silent mutations to alter the sequences recognised by existing siRNAs flanked by complementary sequences. This sequence (shown below) was incorporated into the existing construct by restrictionless cloning based on [89]. The sequence was confirmed by DNA sequencing. Silent mutations introduced are shown below, with nucleotides in bold highlighting substituted nucleotides:

Wild-type *DDX3X* sequence: 5’-CTATATTCCTCCTCATTTA-3’

SiRNA resistant *DDX3X* sequence: 5’-**G**TA**C**AT**C**CC**A**CC**A**CA**CC**T**G**-3

A P2A site and the ORF for superfolder (sf) GFP were obtained by PCR from a pre-existing mammalian co-expression vector by PCR. This was fused to the C-terminus of the existing human DDX3X expression vector by restrictionless cloning. This results in the separate expression of superfolder (sf) GFP in cells transfected with the plasmid in addition to DDX3X.

DDX3X (132-607) was ordered as a codon optimised geneblock from Integrated DNA Technologies including an N-terminal *Bam*HI site, a C-terminal stop codon and a *Not*I site. This fragment was digested and cloned into a *Bam*HI-*Not*I digested and dephosphorylated vector pHEX6P1 (this proprietary vector is based on pGEX6P1 with a 6xHis tag in place of the GST tag).

The full length ORF for human OGT and OGA were cloned into pCMV-FLAG previously as PCR products generated from cDNA. A pre-existing mutant of OGT K842M was sub-cloned from pCMV-FLAG to pCMV-HA to generate a HA tagged inactive OGT protein.

### HEK293T cell culture and transfection

HEK293T cells were cultured in DMEM/F12 medium, supplemented with 10% (v/v) FBS, 1 X GlutaMax, and 5 mL of 5,000 U/mL penicillin/streptomycin. For transfection of CMV-vectors encoding FLAG-tagged constructs, the lipofectamine 3000 transfection kit (ThermoFisher) was used. Briefly, HEK293T cells were seeded at 30% confluency 16 h before transfection, and a 1:2 (w/v) ratio of plasmid to lipofectamine was used in all cases, following the manufacturer’s instructions. Transfections were carried out for 48 h, except for transfections with SiLuciferase, SiDDX3X, SIR WT, or S584A, for which 50,000 cells were transfected and incubated for 72 h.

### Cycloheximide chase assay

HEK293T cells were transfected with FLAG-DDX3X^WT^ or FLAG-DDX3X^Ser584Ala^ as described above. After 48 h, cycloheximide chase was initiated by the addition of 100 μg/mL cycloheximide for the indicated timepoints, after which cells were lysed and analysed by Western blotting.

### Tandem metabolic labelling/SPAAC PEGylation

For investigating endogenous DDX3X O-GlcNAcylation stoichiometry, HEK293T cells were seeded at ∼50% confluence, followed by addition of 200 μM Ac_4_GalNAz or an equal volume of DMSO (vehicle only control) for 16 h. The following day, cells were lysed in RIPA buffer and soluble protein extracted by centrifugation at 13,400 rpm in an Eppendorf mini-spin centrifuge. 200 μg protein was alkylated using iodoacetamide (22.5 mM final concentration; RT, 30 min, in the dark). After alkylation, proteins were chloroform/methanol precipitated by sequential addition of 600 μL MeOH, 150 μL chloroform, and 400 μL water, followed by centrifugation (13,400 x g, 5 min). The upper, aqueous, layer was removed, followed by addition of 450 μL MeOH and centrifugation (13,400 x g, 5 min) to pellet precipitated protein. Precipitated protein was resuspended in 90 μL 25 mM Tris-HCl, pH 8.0, 1% (w/v) SDS, and 10 μL of 10 mM DBCO-mPEG (5 kDa) added to initiate the SPAAC reaction. SPAAC reactions were incubated at 98 °C for 5 min, followed by chloroform/ methanol precipitation and boiling of the precipitated protein in 45 μL 1 x LDS for SDS-PAGE/ Western blotting. For investigating O-GlcNAcylation stoichiometry, HEK293T cells were transfected with FLAG-tagged DDX3X constructs for 48 h as described above. 32 h into the transfection, Ac_4_GalNAz was added to cell media for 16 h, and lysates were PEGylated as described above.

### Recombinant protein expression and purification

A 50 μL aliquot of NiCo21(DE3) *E. coli* was transformed with 1 μL HEXP1 plasmid encoding either His_6_-DDX3X^WT^, His_6_-DDX3X^S584A^. Subsequently, 6 L of LB media (+ 100 μg/mL ampicillin) was inoculated with 10 mL/L of turbid starter culture derived from overnight culture of a single colony in 100 mL LB media (+ ampicillin). Cultures were grown to an OD_600_ of 0.6 at 37 °C (125 rpm), after which expression was induced with 250 μM IPTG. Induced cultures were left overnight at 37 °C (125 rpm). The following day, cultures were spun-down (4,000 x g, 4 °C, 40 min) and bacterial pellets resuspended in a minimal volume of lysis buffer (20 mM HEPES, pH 7.5, 250 mM NaCl, 30 mM imidazole, 0.5 mM TCEP, 1 mM benzamidine, 0.2 mM PMSF, 5 μM leupeptin, ∼0.1 mg/mL lysozyme, and ∼0.1 mg/mL Porcine DNase) and left overnight in a –80 **°**C freezer. The following day, frozen pellets were thawed at room temperature, and lysed via French press. Lysates were spun-down (50,000 x g, 4 **°**C, 1 h) and immediately applied to 10 mL pre-washed Ni^2+^-NTA agarose, followed by incubation for 2 h at 4 °C with rotation. Beads were washed 2 x with lysis buffer, 1 x with lysis buffer containing 1 M NaCl to remove bound nucleic acids, and 7 x more with lysis buffer. His_6_-DDX3X was eluted with 200 mM imidazole. Eluted protein was supplemented with 150 μL of 4 mg/mL PreScission protease for in-solution tag cleavage, and simultaneously dialysed with imidazole free lysis buffer overnight. After negative pulldown with an equal volume of fresh Ni^2+^-NTA beads, the eluent was concentrated to 5 mL and loaded on a HiLoad^TM^ 16/600 Superdex^TM^ 200 size exclusion column with imidazole-free lysis buffer as the mobile phase. Fractions containing protein at the expected Mw were pooled, buffer exchanged into 20 mM HEPES, pH 7.5, 500 mM NaCl, 10% (v/v) glycerol, 0.5 mM TCEP, and concentrated to ∼ 5 mg/mL prior to flash freezing.

### Immunoprecipitation of FLAG-DDX3X

30% confluent HEK293T cells in a 10 cm dish were transfected with 6 μg CMV plasmid encoding FLAG-DDX3X^WT^ or FLAG-DDX3X^S584A^ for 48 h. Cells were lysed in RIPA buffer and 2 mg soluble protein lysate used for FLAG immunoprecipitations. 100 μL of magnetic protein G dynabeads (ThermoFisher) were washed 3 x with PBS followed by incubation with 20 μg mouse anti-FLAG M2 Ab (Sigma Aldrich) for 2 h at 4 **°**C with rotation prior to 3 x PBS washes and resuspension in PBS. Resuspended Ab:protein G immunocomplexes were incubated with 2 mg of transfected HEK293T cell lysate overnight at 4 **°**C, followed by 4 x PBS washes. FLAG-DDX3X was competitively eluted by incubation with 125 μL of 200 μg/mL FLAG peptide. Elutions were immediately used for *in vitro* aggregation experiments as described below.

### In vitro aggregation of immunoprecipitated FLAG-DDX3X

FLAG-DDX3X^WT^ and FLAG-DDX3X^S584A^ were immunoprecipitated from HEK293T lysates as described above. 3 μg of lysozyme and 3 μg rabbit pre-immune IgG were added to 125 μL of immunoprecipitated FLAG-DDX3X as internal standards. Immunoprecipitates were rotated at 800 rpm (37 **°**C) for the indicated time points. At each time point, aggregated protein was pelleted by centrifugation at 13,400 rpm in an Eppendorf mini-spin centrifuge, and boiled in 2 X LDS. A 25 μL aliquot of soluble fraction was mixed with 8 μL 4 X LDS and boiled for analysis by SDS-PAGE. Soluble and pellet fractions were analysed by SDS-PAGE and Coomassie staining.

### Differential scanning fluorimetry

Purified DDX3X^WT^ (132-607) and DDX3X^S584A^ were diluted to 1 mg/mL, and 125 μL of 1 mg/mL DDX3X (132-607) spiked with 1 μL 5000 X SYPRO orange dye. 25 μL of SYPRO orange/ DDX3X mix was aliquoted across 6 wells of a clear PCR plate. A CFX connect Real-Time system was used to measure SYPRO orange fluorescence. After holding at 20 **°**C for 5 seconds, SYPRO orange fluorescence was measured from 20 **°**C to 95 **°**C over 30 min (2.5 **°**C min^-1^). T_m_ values were calculated as described in [90].

### Reverse transcription quantitative PCR (RT-qPCR)

40% confluent HEK293T cells were transfected with FLAG-empty vector or FLAG-OGT^WT^ for 48 h, after which mRNA was extracted using the RNeasy mini-kit (QIAGEN), following manufacturer’s instructions. cDNA synthesis was carried out using 1 μg of mRNA and the qScript cDNA synthesis kit (Quantabio), following manufacturer’s instructions. Reactions consisted of 10 μL PerfeCta SYBR Green FastMix (2 X stock) with 0.3 μL of 10 μM forward and reverse primer each, 4.4 μL milliQ water, and 5 μL 10 ng/μL cDNA. RT-qPCR amplification curves and Cq values were determined using a CFX connect real-time system (cycling conditions: 30 s 95 **°**C, 10 s 95 **°**C, 30 s 60 **°**C; 40 cycles). Primers used are detailed in supplementary table S1. For RT-qPCR of *Ddx3x* in mouse brain samples, total RNA was purified from brain homogenates using the RNAeasy Kit (Qiagen), and then 0.5 to 1 µg of sample RNA was used for reverse transcription with the qScript cDNA Synthesis Kit (Quantabio). All subsequent steps were identical to those described for RT-qPCR of HEK293T mRNA. For mouse RT-qPCR samples were assayed in biological replicates with technical triplicates using the comparative Ct method. The threshold-crossing value was normalized to internal control transcripts (*18S*, *Actb*, and *Pgk1*). Primers used for qPCR analysis are listed in Table S1. Results were normalised to the mean of each corresponding WT replicate set and represented as a fold change relative to WT.

### SiRNA-rescue of cellular proliferation

HEK293T cells were plated at a density of 1 x 10^5^ cells/ well in a 12-well plate and left overnight. Cells were subsequently transfected with 1 μg siRNA against DDX3X (siDDX3X) or Luciferase (siLuciferase), and 0.5 μg plasmid encoding siRNA resistant FLAG-DDX3X^WT^ or FLAG-DDX3X^S584A^. Transfected cells were left for 3 days, after which media was aspirated, followed by 3 x PBS washes to remove dead cells. Cells were subsequently resuspended in 100 μL trypsin, followed by centrifugation at 300 x g (5 min). Trypsin was aspirated and cell pellets were resuspended in 10% (v/v) FBS in PBS, followed by trituration to achieve a homogenous, single cell, suspension. After live cell counting using a LUNA-IITM automated cell counter, cells were spun-down again (300 x g, 5 min) prior to resuspension in lysis buffer.

### Cell cycle analysis

SiRNA rescue of DDX3X was performed as described above. After siRNA rescue, trypsinised cells were centrifuged at 300 x g for 3 min, followed by removal of the supernatant. The cells were then resuspended in 1 mL of PBS with 1% (w/v) FBS. Centrifugation was then repeated at 300 x g for 3 min, and the resulting pellet was resuspended in 0.5 mL of PBS with 1% (w/v) FBS. To fix the cells, 4.5 ml of 70% ethanol at −20 °C was added dropwise during gentle vortexing to maintain a homogenous suspension. The cell suspension was then incubated at 4 °C for 30 min. Following this, the suspension was centrifuged at 300 x g for 3 min, after which the supernatant was aspirated, and cells were rehydrated by the addition of 3 mL PBS supplemented with 1% (w/v) FBS for 2 min. After a further centrifugation at 300 x g for 3 min, the supernatant was removed, and the cells were resuspended in 50 µL of PBS containing 1% (w/v) FBS and 100 µg/mL RNase A for 30 min at room temperature. Subsequently, propidium iodide (PI) was added to a final concentration of 1 µg/mL, followed by incubation in the dark at 4 °C for at least 30 min, preparing the cells for flow cytometry analysis. The cell suspensions were analysed using a NovoCyte 3000 flow cytometer (Agilent, Santa Clara, CA) via a 488 nm laser. The 488/10 detector was used for forward side scatter (FSC) and side scatter (SSC), and the 615/20 detector for PI detection. Data acquisition and analysis were performed using NovoExpress software (version 1.6.2, Agilent, Santa Clara, CA). During data acquisition, multiple gating strategies were employed to isolate single, PI-stained cells for cell cycle analysis. Initially, cells were gated based on size and granularity using FSC area (FSC-A) and SSC area (SSC-A). These gated cells were then analysed by comparing FSC height (FSC-H) to FSC-A to identify single cells. The single cells were further plotted on a PI area (PI-A) versus FSC-A graph to select PI-stained cells. Finally, these gated cells were subjected to cell cycle analysis using the built-in function of the software, employing the Watson Pragmatic model for data fitting.

### MTT assay

To identify the linear range in which the number of viable cells is directly proportional to formazan absorbance at 560 nm, a standard curve was generated by plating 2.5 x 10^4^ to 2.5 x 10^5^ HEK293T cells in a 12-well plate. After overnight incubation, cell culture media was removed and 1 mL of 0.5 mg/mL thiazolyl blue tetrazolium bromide, dissolved in media, was added to each well. Cells were incubated for 4 h prior to removal of the cell culture medium and addition of 100 μL DMSO for 5 min to solubilise formazan crystals. DMSO-solubilised formazan was transferred to a 96-well plate and absorbance at 560 nm measured using a SpectraMax iD5. To ensure cell viability counts were within the linear range of detection, siRNA rescue experiments were performed using 1 x 10^5^ cells. After 72 h, the MTT assay was repeated as described above, with the additional step of diluting the DMSO-solubilised formazan crystals 1:16 in DMSO. 100 μL of the diluted formazan crystals were transferred to a 96-well plate and absorbance measured at 560 nm as described above.

### Western blotting

40 μg of PEGylated glycoprotein or 20 μg of cell lysate were run on a 4-12% NuPAGE Bis-Tris gel for 2 h (120 V). Proteins were transferred onto nitrocellulose membrane using an Invitrogen semi-dry transfer system for 1 h (20 V). After blocking in 5% (w/v) bovine serum albumin dissolved in TBS (+ 0.1% (v/v) Tween-20) for 30 min (RT) membranes were incubated with the indicated antibodies for 1-2 h at RT, followed by 3 x 5 min TBST washes. Membranes were then incubated with the corresponding secondary antibody for 1 h (RT), prior to imaging using a Li-COR Odyssey.

### Statistics

Western blot band signals and Coomassie blue staining intensities were integrated using ImageStudioLite (v 5.2.5) and statistical analyses carried out in GraphPad Prism (v 9.0). All statistical analyses pertaining to the BrainCloud atlas was performed in RStudio (see above for details). Sigmoidal, monophasic, curves were extracted from DSF data and analysed in GraphPad Prism (v 9.0). Throughout this manuscript, ns: not significant, *: *P* < 0.05, **: *P* < 0.01, ***: *P* < 0.001, **** *P* < 0.0001.

### Animal husbandry

Animals were housed in ventilated cages with water and food available *ad libitum* and 12/12 h light/dark cycles. All animal studies and breeding were performed on in accordance with the Animal (Scientific Procedures) Act of 1986 for the care and use of laboratory animals. Procedures were carried under United Kingdom Home Office Regulation (Personal Project Licence PP8833203) with approval by the Welfare and Ethical Use of Animals Committee of University of Dundee.

### Tissue collection and disruption

Brain tissues from 80-91 (OGT^C921Y^), 86-90 (OGT^N648Y^) and 132-134 (OGT^N657K^) days old male mice were rapidly dissected, snap frozen in liquid nitrogen, and stored at −80 °C. Tissues were disrupted in Phosphate-Buffered Saline two times at 5000 rpm for 30 s with 10 s break using Precellys^®^ 24 Touch homogenizer (Bertin Technologies). Homogenates were split in half for further protein and RNA extractions.

## Acknowledgements

This work was funded by a Novo Nordisk Fonden Laureate award (NNF21OC0065969) and a Villum Fonden Investigator (00054496) to D.M.F.v.A. C.W.M was funded by the BBSRC EastBio Doctoral Training Programme.

## Author contributions

C.W.M. and D.M.F.v.A. conceived the study; C.W.M, H.Y, and F.A. performed experiments; M.S-J performed bioinformatic analysis of frontal cortex transcriptomes; A.M.D’A analysed clinical literature for DDX3X ID and OGT-CDG; C.W.M., F.A., H.Y., M.S-J, A.M.D’A and D.M.F.v.A. analysed data; C.W.M. and D.M.F.v.A. wrote the manuscript with input from all authors.

## Supplementary figure legends

**Supplementary figure S1:**
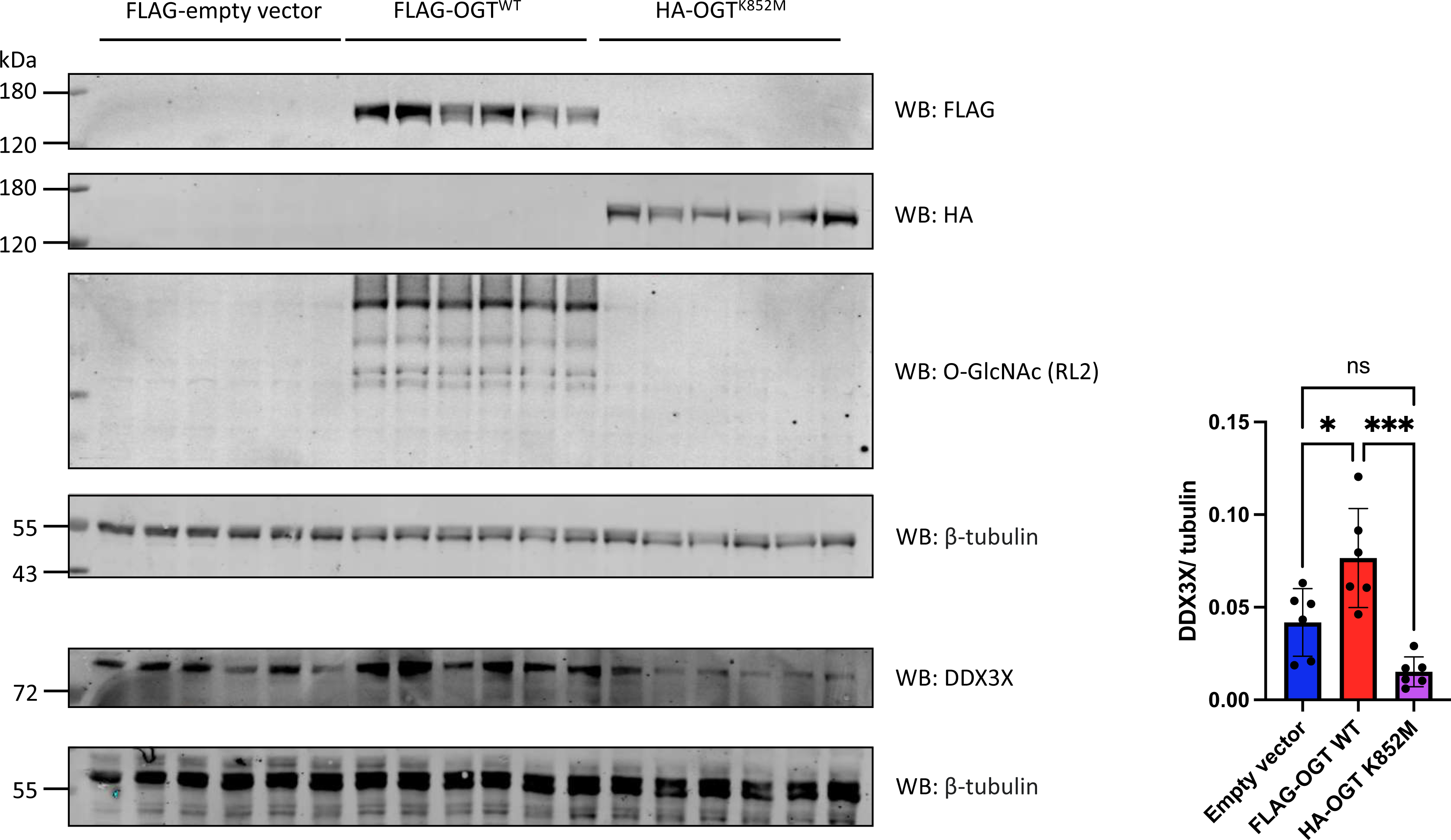
The glycosyltransferase activity of OGT is required for increased DDX3X protein levels in HEK293T cells following OGT transfection. HEK293T cells were transiently transfected with FLAG-empty vector, N-terminally FLAG-tagged OGT^WT^, or N-terminally HA-tagged OGT^K852M^. Total O-GlcNAc levels were measured relative to β-tubulin by blotting with the pan-specific O-GlcNAc antibody RL2 as a control for OGT^WT^ activity and verification of the catalytic inactivity of OGT^K852M^. DDX3X levels were also measured relative to β-tubulin. Normalised DDX3X levels are plotted as the mean +/- S.E.M. Data were analysed using a one-way ANOVA (N = 6 technical replicates). ns: not significant; *: *P*<0.05; ***: *P*<0.001.

**Supplementary figure S2:**
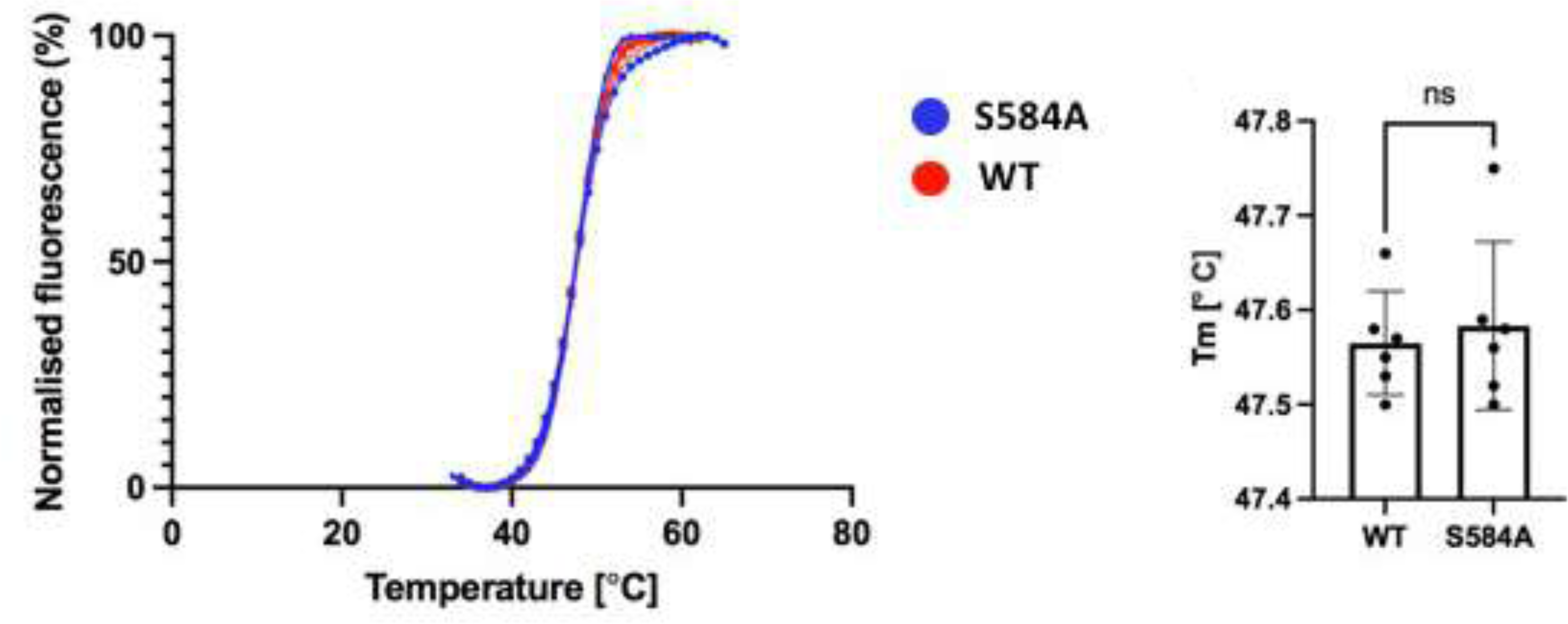
Alanine mutagenesis of Ser584 does not affect the thermal stability of recombinant DDX3X (132-607). Soluble truncates of DDX3X^WT^ and DDX3X^S584A^ (residues 132-607) were expressed in *E. coli* as N-terminal His_6_ tag fusions and purified (see materials and methods for details regarding purification). 1mg/ mL of DDX3X^WT^ and DDX3X^S584A^ were supplemented with SYPRO orange and thermal denaturation curves measured. The extracted monophasic transitions were extracted from the fluorescence measurements and are plotted. Additionally, the T_m_ values for DDX3X^WT^ and DDX3X^S584A^ are plotted +/- S.E.M. N = 6 technical replicates. Data were analysed using Student’s unpaired *t*-test. ns: not significant.

**Supplementary figure S3:**
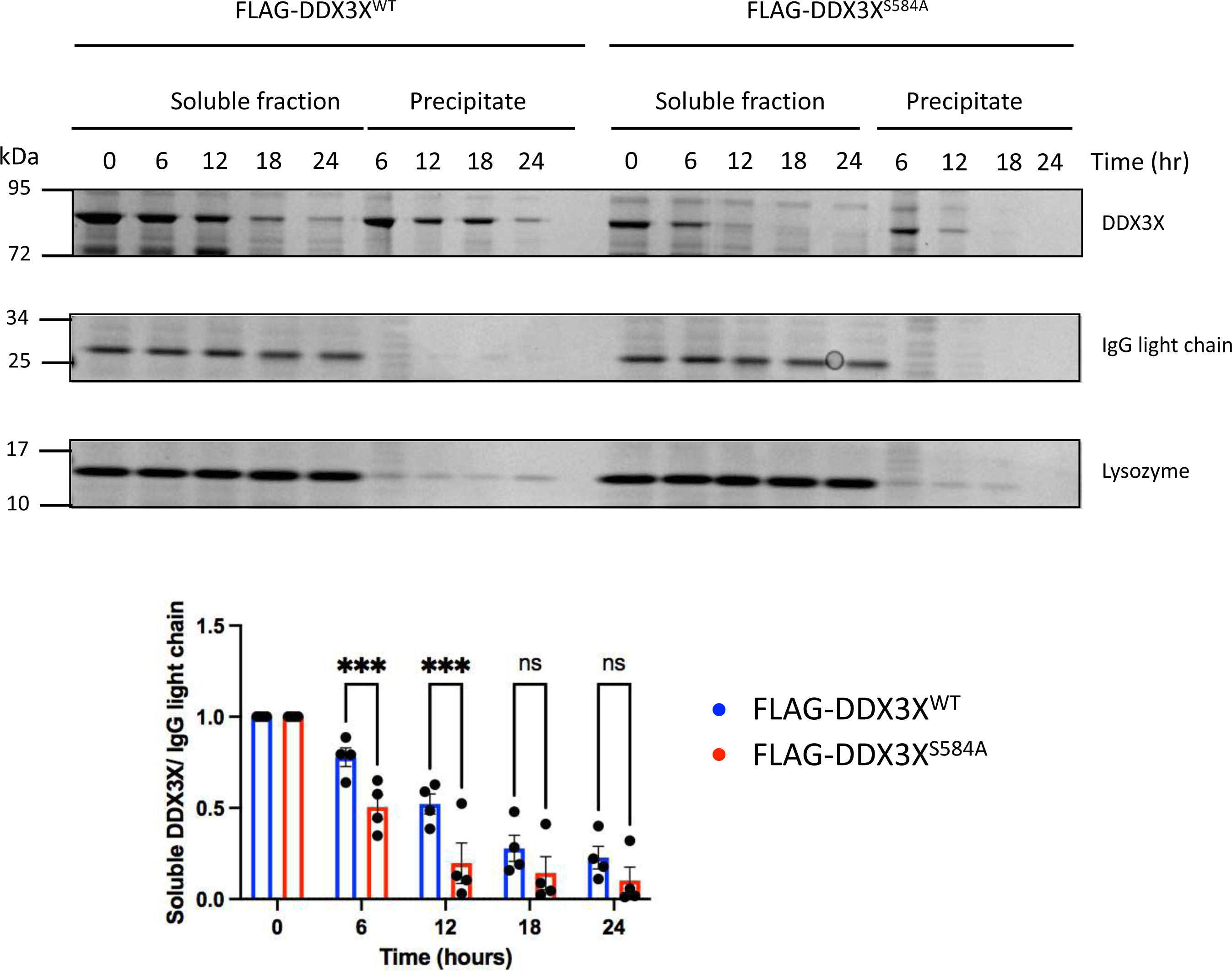
*In vitro* aggregation of immunoprecipitated FLAG-DDX3X^WT^ and FLAG-DDX3X^S584A^. FLAG-DDX3X^WT^ and FLAG-DDX3X^S584A^ were immunoprecipitated from transfected HEK293T lysate via the N-terminal FLAG tag. Immunoprecipitates were spiked with 3 μg of lysozyme and 3 μg rabbit pre-immune IgG prior to incubate at 37 °C for 24 h. At each time point (0,6,12,18,24), immunoprecipitates were centrifuged to remove insoluble protein aggregates. Soluble and insoluble fractions for each timepoint were analysed by SDS-PAGE and Coomassie staining. Soluble DDX3X levels were measured via band densitometry and normalised to the rabbit pre-immune IgG light chain. Data were analysed using a two-way ANOVA (N = 4 independent experiments), and plotted as the mean +/- S.E.M. ns: not significant; ***:*P*<0.001.

**Supplementary figure S4:**
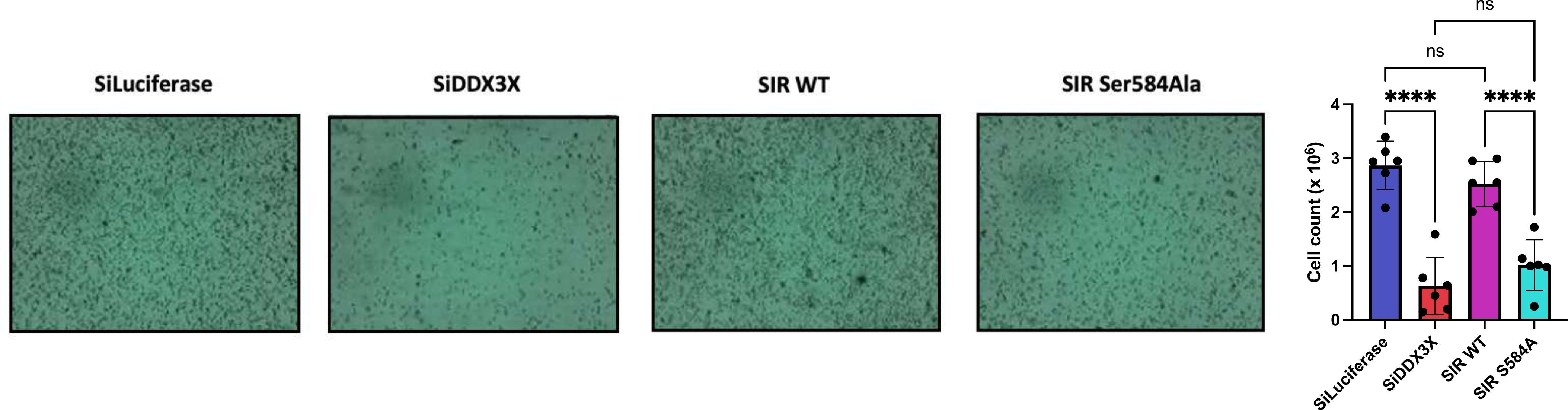
representative images of live cells after siRNA knockdown of *DDX3X* and rescue with co-transfected siRNA-resistant DDX3X^WT^ and DDX3X^S584A^. Representative images of live cells after knockdown of endogenous *DDX3X* and rescue with co-transfected siRNA-resistant FLAG-DDX3X^WT^ or FLAG-DDX3X^S584A^. PBS washes were used to remove apoptosed or necroptosed cells and the remaining live cells were stained with trypan blue. Live cell counts were calculated using a LUNA-IITM live cell counter. Live cell counts across 6 technical replicates are plotted as the mean +/- 1 SD. Data were analysed using a one-way ANOVA. ns: not significant; ****:*P*<0.0001.

**Supplementary table S1.**
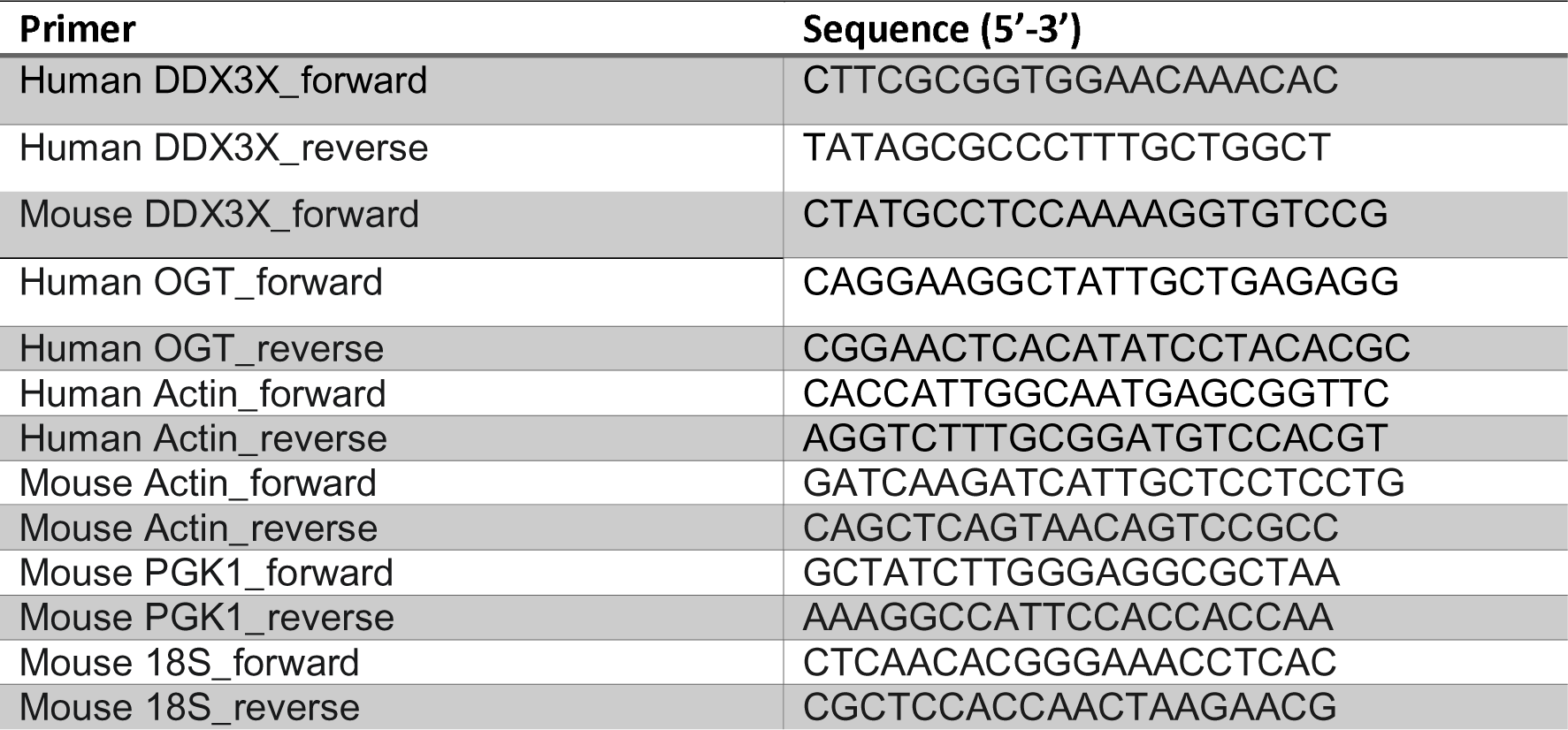
Primers used in this study for RT-qPCR.

